# The Zip4 protein directly couples meiotic crossover formation to synaptonemal complex assembly

**DOI:** 10.1101/2021.08.13.456249

**Authors:** Alexandra Pyatnitskaya, Jessica Andreani, Raphaël Guérois, Arnaud De Muyt, Valérie Borde

## Abstract

Meiotic recombination is triggered by programmed double-strand breaks (DSBs), a subset of these being repaired as crossovers, promoted by eight evolutionarily conserved proteins, named ZMM. Crossover formation is functionally linked to synaptonemal complex (SC) assembly between homologous chromosomes, but the underlying mechanism is unknown. Here we show that Ecm11, a SC central element protein, localizes on both DSB sites and sites that attach chromatin loops to the chromosome axis, which are the starting points of SC formation, in a way that strictly requires the ZMM protein Zip4. Furthermore, Zip4 directly interacts with Ecm11 and point mutants that specifically abolish this interaction lose Ecm11 binding to chromosomes and exhibit defective SC assembly. This can be partially rescued by artificially tethering interaction-defective Ecm11 to Zip4. Mechanistically, this direct connection ensuring SC assembly from CO sites could be a way for the meiotic cell to shut down further DSB formation once enough recombination sites have been selected for crossovers, thereby preventing excess crossovers. Finally, the mammalian ortholog of Zip4, TEX11, also interacts with the SC central element TEX12, suggesting a general mechanism.

## Introduction

Meiosis is a highly conserved process among organisms with sexual development. It produces four haploid gametes from one diploid cell by executing two successive rounds of cell division preceding one round of DNA replication (Hunter, 2015). A unique defining feature of meiosis is the pairing/synapsis and homologous recombination between parental chromosomes (homologs). Recombination is initiated by programmed DNA double-strand break (DSB) formation by the topoisomerase-related Spo11 protein together with several meiotic protein partners (Yadav and Claeys Bouuaert, 2021). Following DSB formation, the combined action of endo- and exonucleases leads to resection of the DSBs 5’ ends, creating 3’ single-strand DNA tails. The strand exchange proteins Rad51 and Dmc1 bind to these tails, and form a nucleofilament that invades the homologous chromosome. This results in the formation of a D-loop intermediate that goes through various steps of maturation, leading to two possible outcomes: a crossover (CO) with a physical exchange between chromosomal arms, or a non-crossover (NCO). Meiotic COs can be subdivided in two classes, with class I COs representing ∼85 % of total COs formed in budding yeast, mammals and plants. A characteristic of class I COs is that they are more evenly spaced from each other than would be expected from a random distribution, phenomenon referred to as “interference” (Berchowitz and Copenhaver, 2010). The ZMM group of proteins (for Zip1-4, Msh4-5, Mer3, Spo16) is the major actor promoting class I CO formation (Börner et al., 2004; Pyatnitskaya et al., 2019). Molecularly, these proteins are proposed to act on D-loop recombination intermediates by protecting them against their dismantling by helicases, which would lead to NCO (De Muyt et al., 2012; Zakharyevich et al., 2012). ZMM-protected intermediates are then maturated into a particular DNA structure that will be further processed into CO by the endonuclease activity of the MutLγ (Mlh1-Mlh3)-Exo1 complex (De Muyt et al., 2012; Hunter and Kleckner, 2001; Zakharyevich et al., 2012). Among the ZMM proteins, the Zip2-Zip4- Spo16 complex plays a predominant role, through its XPF-ERCC1-like module, in specifically binding branched recombination intermediates (Arora and Corbett, 2019; De Muyt et al., 2018). In addition, this complex has a scaffolding activity through its Zip4 subunit. Indeed, Zip4 interacts with several other ZMM proteins as well as with Red1, a component of the meiotic chromosome axis (axial element), forming the lateral element of the synaptonemal complex (SC) during homolog synapsis (De Muyt et al., 2018). The SC appears concomitantly with the maturation of the ZMM-protected recombination intermediates. It is composed of two lateral elements physically maintained together at a precise distance of 100 nm by a central region (Zickler and Kleckner, 1999). SC assembly begins with the formation of the axial element along each pair of sister chromatids. Polymerization of axial elements leads to arrays of chromatin loops tethered at their bases to the axial proteins, among which the meiosis-specific Hop1 and Red1 proteins, and cohesin containing the Rec8 subunit (Klein et al., 1999; Panizza et al., 2011; Smith and Roeder, 1997). Homologous chromosomes co-align across their length, then, the central region polymerizes from punctuate sites to progressively connect axial elements of the two homologs until the chromosomes are synapsed along their entire length (Boer and Heyting, 2006; Moses, 1969). In budding yeast, the central region is composed of the transverse filament Zip1 and the central element, including Ecm11 and Gmc2, which facilitate Zip1 assembly (Gao and Colaiácovo, 2018; Humphryes et al., 2013; Sym et al., 1993).

In budding yeast, CO formation and SC polymerization are spatially and functionally related. Indeed, SC polymerization often initiates from sites called SICs (for “Synapsis Initiation Complex”), enriched in ZMM, and therefore likely representing recombination intermediates, where ZMM were shown to bind by ChIP-seq approaches (Agarwal and Roeder, 2000; Chua and Roeder, 1998; De Muyt et al., 2018; Serrentino et al., 2013; Shinohara et al., 2008; Tsubouchi et al., 2006). In *Sordaria macrospora*, SC nucleates and emanates from one side of recombination nodules, structures that are particularly dense on electron microscopy images and are predicted to be aggregates of active recombination proteins including ZMMs (Dubois et al., 2019). In mammals, whether SC polymerization starts from ZMM-enriched sites is still not fully established. However, a large majority of RNF212, related to the ZMM Zip3 protein, colocalizes with initial stretches of SYCP1, the mouse homolog of Zip1, suggesting that such mechanism occurs in mammals (Reynolds et al., 2013). Moreover, the absence of ZMM proteins leads to synapsis defects in both budding yeast and *Sordaria*, suggesting that stabilization of CO precursors is important for correct SC polymerization (Agarwal and Roeder, 2000; Chua and Roeder, 1998; Dubois et al., 2019; Espagne et al., 2011; Shinohara et al., 2008; Storlazzi et al., 2010; Tsubouchi et al., 2006). Similarly, several mouse ZMM mutants (*Msh4^-/-^*, *Msh5^-/-^*, *Hfm1/Mer3^-/-^*, *Shoc1/Zip2^-/-^*) show strong synapsis defects (reviewed in (Pyatnitskaya et al., 2019)). On the other hand, the SC is involved in crossover formation. Whether the SC is involved in mediating crossover interference has been investigated in several model organisms. In budding yeast, this is clearly not the case. A deletion mutant of Zip1, the transverse filament of the SC but also a ZMM protein, is defective in interfering COs. However, in mutants where Zip1 still binds recombination intermediates but does not polymerize, such as the Nter deletion *zip1N1* mutant or the central element *ecm11*Δ and *gmc2*Δ mutants, CO still interfere, although the strength of interference is slightly, but significantly reduced (Lee et al., 2021; Voelkel-Meiman et al., 2015, 2016). These mutants likely preserve the Zip1 “ZMM function” intact, which is independent of its SC assembly function (Börner et al., 2004; Chen et al., 2015; Voelkel-Meiman et al., 2015, 2016). Although SC polymerization is not formally required for the formation of interfering COs, it does seem to play a regulatory role in their distribution. In budding yeast, despite wild-type spore viability, *zip1N1*, *ecm11*Δ and *gmc2*Δ mutants show increased CO frequency on certain chromosomes, suggesting that the SC could limit ZMM-dependent CO formation (Lee et al., 2021; Voelkel-Meiman et al., 2016, 2019). This may be explained at least in part by recent findings that Ecm11- and Gmc2- dependent SC assembly downregulates DSB formation by Spo11 (Lee et al., 2021; Mu et al., 2020). Similarly, in plants, mutants of the transverse filament *ZEP1 and AtZYP1* in rice and *Arabidopsis*, respectively, show more COs, indicating that like in budding yeast, the SC is regulating crossover frequencies. However, contrary to budding yeast, these crossovers lost interference although they still seem to depend on ZMM (Capilla-Pérez et al., 2021; France et al., 2021; Wang et al., 2010). Similarly in *C.elegans*, partial depletion of the synaptonemal complex central region proteins reduces the effective distance over which interference operates, suggesting that synaptonemal complex proteins also limit crossovers in nematode (Libuda et al., 2013). These apparent differences with fungi deserve further investigation but may stem from the fact that progression through meiosis in plants is not affected by the absence of ZMM proteins.

Despite the temporal and spatial relationships between CO formation and SC assembly, the underlying physical connections between the two processes are elusive. Here, we uncover a direct interaction between the ZMM protein Zip4 and the central components of the SC Ecm11 and Gmc2, which is essential for the recruitment of the Ecm11 protein to chromosomes and consequently for SC polymerization. We propose a model in which Zip4 brings Ecm11 to recombination sites that are prone to form Cos and helps the transverse filament protein Zip1 to nucleate from this location, ensuring a control of recombination starting locally from sites engaged in the crossover repair pathway.

## Results

### The central element protein Ecm11 interacts with Zip4 and is recruited to DSB and axis-attachment sites

To investigate possible physical connections between crossover formation and synaptonemal complex assembly pathways, we systematically tested by yeast two-hybrid the interactions between ZMM proteins and the known SC components (Fig. 1A). The only interactions were between Zip4 and each of the two known SC central elements, Ecm11 and Gmc2 (Fig. 1A). We confirmed that the endogenous proteins interact in meiotic cells, by coimmunoprecipitating Zip4-Flag protein with Ecm11-TAP (Fig. 1B). Since Zip4 is known to be recruited to recombination sites, we next asked if Ecm11 shows a similar binding pattern by mapping Ecm11 binding sites at 5 h in meiosis, the expected time of recombination (Hunter and Kleckner, 2001), using spike-in calibrated ChIP-seq (Fig. 1C-D) (Hu et al., 2015). Strikingly, Ecm11 preferentially localized, like Zip4, at DSB hotspots, indicating that Ecm11 is present at the recombination intermediates. In addition, Ecm11 also preferentially associated to Red1 binding sites, which define the basis of chromatin loops attached to the chromosome axis, where SC polymerizes, consistent with Ecm11 being a component of the SC (Fig. 1C-D). Looking at the kinetics of Ecm11 association with chromatin by ChIP-qPCR revealed that Ecm11 binding to DSB and axis-attachment sites was maximum at 4 - 5 h in meiosis, during recombination (Fig. 1E). Then, we sought to find the determinants for Ecm11 association to chromosomes, first by testing if Zip4 is involved. Indeed, Ecm11 recruitment to chromatin was drastically reduced in a *zip4*Δ mutant on both DSB and axis sites (Fig. 1C-E). Previous studies have suggested that Zip1 may be important for Ecm11 loading (Voelkel-Meiman et al., 2015, 2016). Interestingly, the recruitment of Ecm11 to DSB hotspots was only partially reduced in absence of Zip1, while the association with the axis-binding sites was more strongly impaired (Fig. 1C-E, *zip1*Δ). We asked if the reduced Ecm11 association to chromosomes in *zip1*Δ may be a consequence of reduced Zip4 binding to chromosomes. Indeed, Zip4 enrichment was strongly reduced in absence of Zip1, which is likely the reason for reduced Ecm11 binding in *zip1*Δ (Supplemental Fig. S1). Zip1 therefore seems important for full Ecm11 localization at the SC, likely because Ecm11-Gmc2 co-polymerize together with Zip1, but less so for its recruitment to recombination sites.

**Figure 1:**
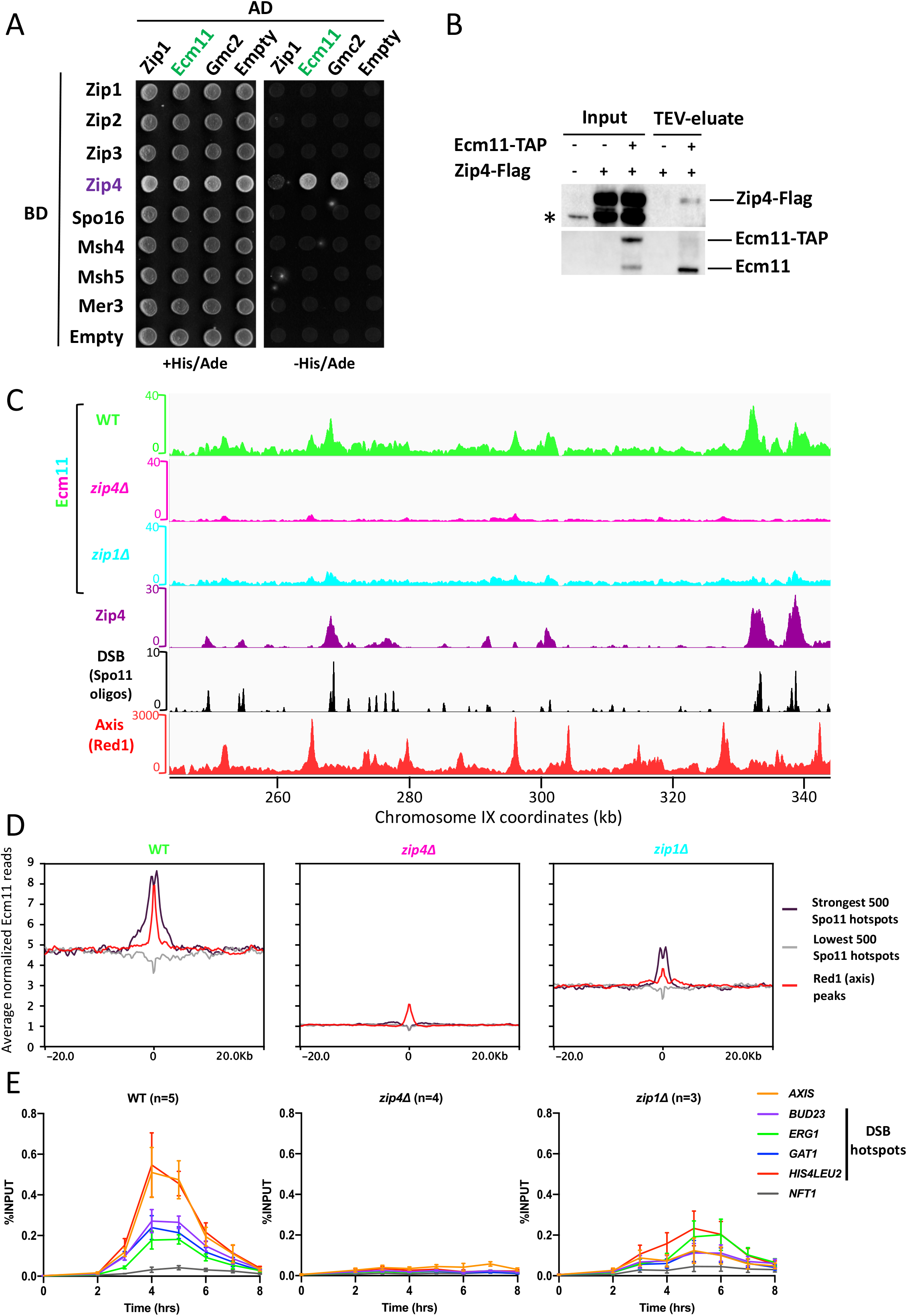
Ecm11 localization on DSBs and axis-attachment sites is dependent on Zip4. A. Yeast two-hybrid interaction analysis between SC components Ecm11 and Gmc2 and the ZMM proteins. Prey and baits are fused with the GAL4 Activation Domain (GAL4-AD) and with the GAL4 DNA-Binding Domain (GAL4-BD), respectively. Interaction results in growth on the selective –His/Ade medium. B. Co-immunoprecipitation between Zip4-Flag and Ecm11-TAP from meiotic cells at 5 h in meiosis, analyzed by western blot. The asterisk indicates a non-specific cross-reacting band and possible products of Zip4-Flag degradation. C. ChIP-seq DNA-binding of Ecm11-Flag in *WT*, *zip4*Δ and *zip1*Δ strains. Normalized data are smoothed with a 200-bp window. Zip4-binding profile is also shown (De Muyt et al., 2018). DSB sites are mapped by Spo11 oligos (Zhu and Keeney, 2015) and axis-attachment sites by Red1 binding profile (Sun et al., 2015). D. Average Ecm11 ChIP-seq signal of data shown in A. at the indicated features. Alignments were performed on the Spo11 hotspots midpoints from (Zhu and Keeney, 2015) and Red1 peaks summits from (Sun et al., 2015). E. ChIP monitoring of Ecm11-Flag association with different chromosomal regions, measured by qPCR using primers that cover the indicated regions. Same strains as in C. are used. Values are the mean ± SEM of the indicated number of independent experiments.

Finally, our quantitative Ecm11 ChIP-seq data also revealed relatively uniform Ecm11 binding outside of recombination hotspots and axis sites, which was strongly diminished in the absence of Zip4 (Fig. 1C,D). This was confirmed by qPCR with the enrichment of Ecm11 at the *NFT1* site, a locus that shows neither DSB nor detectable axis protein signal (Fig. 1E) (Sun et al., 2015; Zhu and Keeney, 2015). Such random binding may reflect, in addition to preferential sites, a mobility of the loop-attachment sites to the chromosome axis, that may be mediated by constant loop extrusion by cohesin at the basis of these loops, as recently shown in mammalian cells (Fudenberg et al., 2016).

Altogether, we conclude that Ecm11 localizes at recombination sites and along the chromosome axis, in a Zip4-dependent manner.

### Zip4-Ecm11 interaction is important for normal SC polymerization

To further investigate the role of the Zip4-Ecm11 interaction in meiosis, we characterized the domains of Zip4 and Ecm11 mediating the interaction. Zip4 encompasses 21 TPR (TetratricoPeptide Repeat) motifs spanning the whole length of Zip4 and ends with a C-terminal alpha-helix (Perry et al., 2005) (Fig. 2A). We generated a reliable 3D model of Zip4, which revealed an extensive surface featuring four distinct conserved patches likely to be involved in protein interactions (Fig. 2B). We used this model to delineate fragments of Zip4, sufficiently long to enable proper folding and maintain interactions without disrupting the conserved patches (Fig. 2A). Yeast two-hybrid experiments showed that Ecm11 interacts with the last C-ter fragment that contains the most conserved patch of Zip4 (Fig. 2A and Supplemental Fig. S2A). A search for conserved and surface-exposed aminoacids potentially involved in protein-protein interactions in this region uncovered a highly conserved aromatic-asparagine motif (residues W918-N919 in *S. cerevisiae* Zip4) (Fig. 2C). This motif is often present in different binding scaffolds, such as in the Armadillo repeats of importin *α* for interaction with NLS motifs (Fontes et al., 2000). Exposed and conserved asparagine residues in these domains are typically found to mediate specific interactions with the backbone amide groups of the binding partner. Therefore, we mutated this motif by substituting the asparagine 919 with a glutamine (Zip4N919Q), changing only the steric hindrance to have a minimal effect on the rest of the protein. Remarkably, Zip4N919Q completely lost its interaction with Ecm11, as assessed by yeast two-hybrid, while keeping its interaction with Zip4’s other known partners Zip3 and Zip2 (Fig. 2A and Supplemental Fig. S2B). Co-IP experiments from meiotic cells also confirmed that the interaction between Zip4N919Q mutant and Ecm11 was strongly reduced *in vivo* (Fig. 2D).

**Figure 2:**
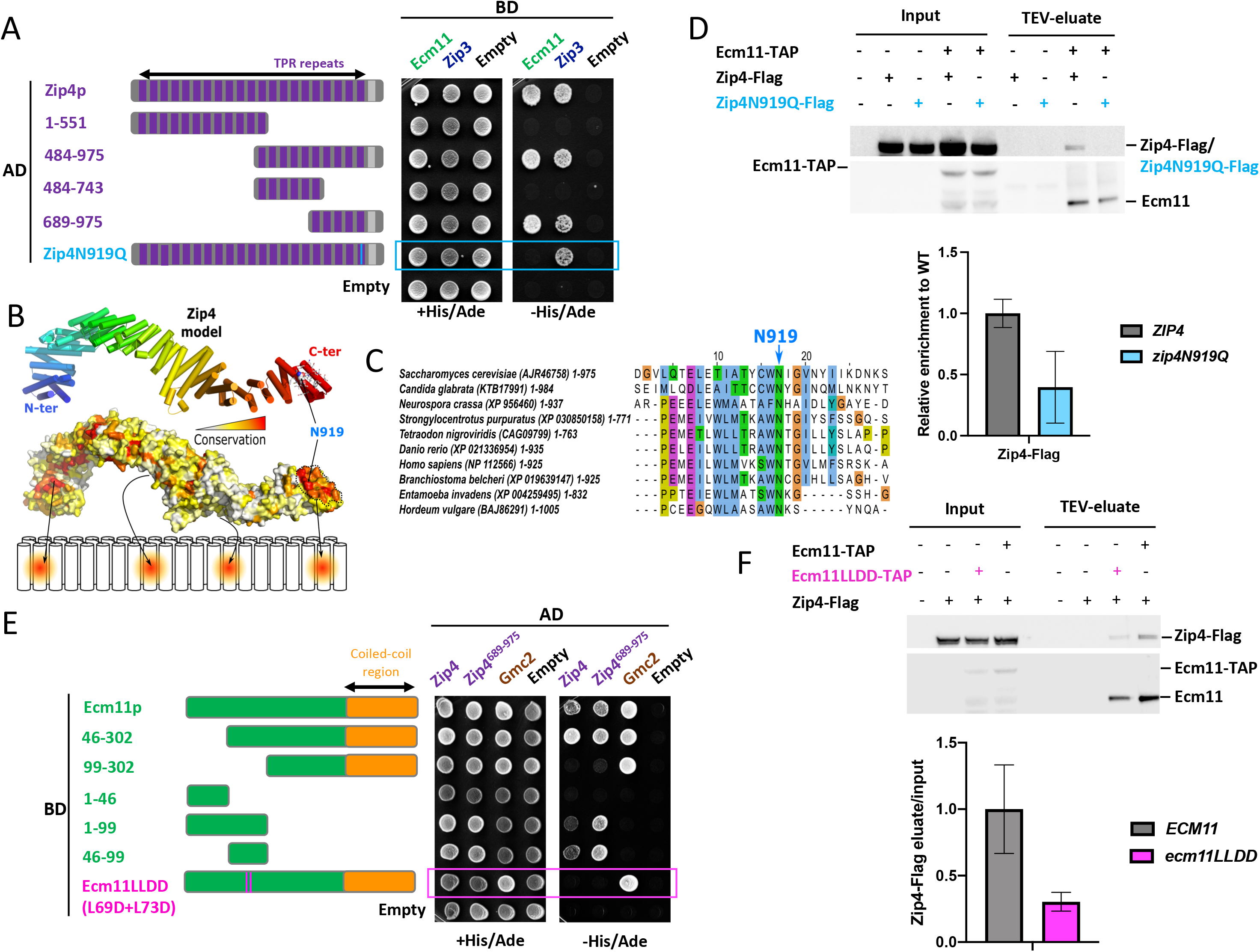
Zip4 specifically interacts with Ecm11. **A.** Delineation of the Ecm11-interacting domain in Zip4 by two-hybrid assays. Indicated fragments of Zip4 were fused to GAL4-AD and tested in combination with a GAL4-BD-Ecm11 or -Zip3 fusion. The blue frame indicates the absence of interaction between Zip4N919Q and Ecm11. **B.** 3D model of Zip4 TPR revealing 4 conserved surface patches. The degree of conservation is shown. **C.** Alignment of Zip4 C-terminal TPR domain. **D.** Co-immunoprecipitation between Zip4-Flag, Zip4N919Q-Flag and Ecm11-TAP from meiotic cells at 5 h in meiosis, analysed by western blot. Levels of Zip4-Flag and Zip4N919Q-Flag were quantified relative to the input and normalized by Ecm11-TAP levels. Values are the mean ± SD of two independent experiments. **E.** Same assay as in A. Ecm11 domains were fused to GAL4-AD and tested in combination with GAL4-BD-Zip4 or GAL4-BD-Zip4-689-971. The pink frame indicates the loss of interaction between Ecm11LLDD and Zip4. **F.** Co-immunoprecipitation of Zip4-Flag with Ecm11-TAP or with Ecm11LLDD-TAP from meiotic cells at 4 h in meiosis, analysed by western blot. Levels of Zip4-Flag coimmunoprecipitated with Ecm11-TAP or with Ecm11LLDD-TAP were quantified relative to the input and normalized by Ecm11-TAP or Ecm11LLDD-TAP levels. Values are the mean ± SD of two independent experiments.

To delineate the Ecm11 regions interacting with either Zip4, we further analyzed the C-terminal conserved patch of Zip4 in the vicinity of asparagine 919 and identified a set of four exposed apolar residues distributed over the 19^th^ and 20^th^ TPR repeats (Supplemental Fig. S3A), suggesting that the Ecm11 binding region should contain a significant number of conserved hydrophobic residues to interact with this region. From the multiple sequence alignment of Ecm11 (Supplemental Fig. S3B), sequence analysis predicts the existence of a long disordered N-terminal tail extended by a 70-residue coiled-coil in the C-terminus. A short stretch spanning residues 68-76 in the disordered tail contains two conserved and hydrophobic positions and a propensity to adopt a helical conformation, making this region a good candidate for interacting with Zip4 in the vicinity of N919. For the interaction between Ecm11 and Gmc2, we exploited a coevolution-based analysis, which suggested that the C-terminal coiled-coil of Ecm11 could most likely form anti-parallel and parallel coiled-coils with Gmc2 (Supplemental Fig. S7). We validated these predictions by Y2H experiments, where the domain 46-99 of Ecm11 was sufficient to interact with Zip4 while the coiled coil region 212-302 was critical for Gmc2 interaction but not for Zip4 binding (Fig. 2E and supplemental Fig. S3C). Within the 46-99 region of Ecm11, two well-conserved hydrophobic residues, leucines L69 and L73, are good candidates for Zip4 interaction (Supplemental Fig. S3B). Indeed, their mutation to aspartate (generating the Ecm11L69D-L73D mutant, hereafter called Ecm11LLDD) disrupted the Ecm11-Zip4 interaction, while preserving the Ecm11-Gmc2 interaction in yeast two-hybrid (Fig. 2E). This effect was confirmed *in vivo* where the interaction of Ecm11LLDD with Zip4 was decreased (Fig. 2F). Altogether, these results indicate that Zip4 and Ecm11 interact directly, through a region on Ecm11 distinct from the Gmc2-binding region, which establishes a physical connection between CO formation and SC assembly processes.

### Disturbing Zip4-Ecm11 interaction strongly affects Ecm11 recruitment and SC assembly

To address the function of the interaction between Zip4 and Ecm11 during meiotic prophase I, we first assessed spore viability and meiotic progression in the interaction mutants. The *zip4N919Q* and *ecm11LLD* mutants showed wild-type spore viability, like *ecm11*Δ but in sharp contrast with *zip4*Δ (Fig. 3A). In addition, they both showed a shorter delay in meiotic divisions (3 h and 1.5 h, respectively) than *zip4*Δ (more than 5 h) (Supplemental Fig. S4A). Together, these data suggest that the Zip4-Ecm11 interaction is not needed for Zip4 ZMM functions in CO formation. We noted that the *zip4N919Q* was slightly more delayed than *ecm11*Δ (1.5 h delay), which may be related to the lower levels of the Zip4^N919Q^ protein detected during meiosis (Supplemental Fig. S4B). The Zip4 WN motif exhibits a degree of conservation from yeast to human much higher than that of Ecm11 whose homologs are only found in fungi (Fig. 2C and Supplemental Fig. S3B). Therefore, we cannot exclude that the WN motif has additional functions, besides interaction with Ecm11, such as interaction with a chaperone, that would ensure Zip4 stability. However, since it would involve the same residues as for Ecm11 interaction, this would occur at a different step, such as during Zip4 “ZMM” activities.

**Figure 3:**
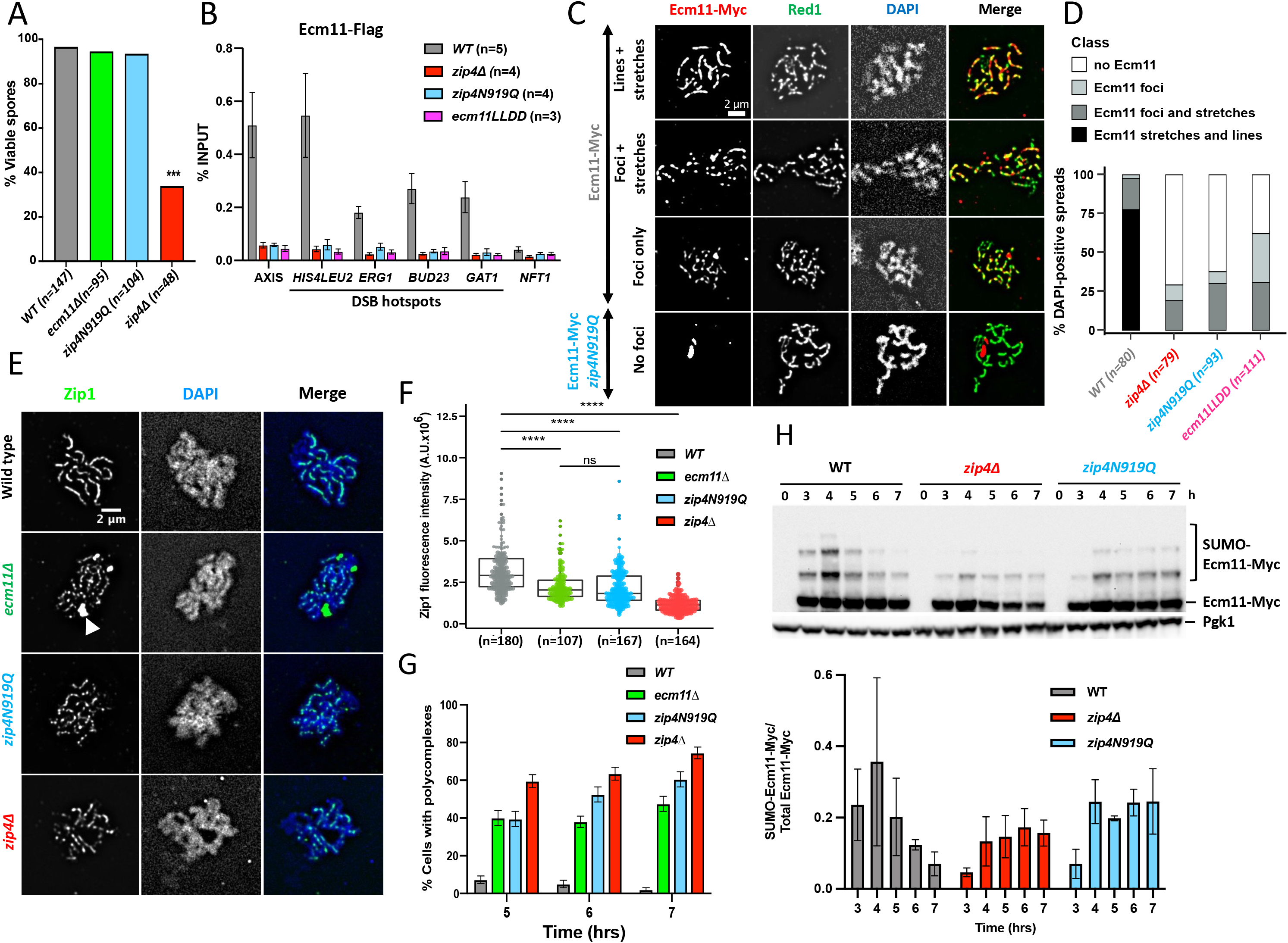
Synaptonemal complex assembly depends on the interaction of Ecm11 with Zip4. A. Spore viability assays of strains with the indicated genotype. Numbers of dissected tetrads are indicated. ****p < 0.0001, Fisher’s exact test. B. Maximum levels of Ecm11-Flag or Ecm11LLDD-Flag in the indicated strains measured by quantitative PCR (qPCR) using primers that cover the indicated regions are shown. Values are the mean ± SEM from at least three independent experiments. The full corresponding time courses are in Fig. 1D and Supplemental Fig. S4. C. Ecm11-Myc localization on surface-spread chromosomes in the indicated strains. Red: anti-Myc (red); green: anti-Red1, blue: DAPI. Red1-positive spreads were divided in four categories: 1) exhibiting stretches and lines of Ecm11 – synapsis almost complete or complete, 2) exhibiting foci and stretches of Ecm11 – partial synapsis, 3) exhibiting only Ecm11 foci – dotty pattern, 4) exhibiting no Ecm11. Representative pictures are shown for the indicated strain. The pictures for the other strains are in Supplemental Fig. S5. D. Quantification of the classes shown in C. The number of counted spreads is indicated. E. Zip1 localization on surface-spread chromosomes in the indicated strains. Only pachytene or pachytene-like stages were considered. Green: anti-Zip1; blue: DAPI (DNA). White arrow: Zip1 polycomplex. F. Quantification of Zip1 intensity observed in E. Numbers of spreads are indicated for each genotype. ****: p-value<0.0001, Wilcoxon test. G. Quantification of DAPI-positive spreads showing a polycomplex. At least 200 spreads were considered for each condition. Values are % cells ± SD of the proportion.H. Ecm11 SUMOylation in the indicated strains analyzed by western blot. Quantification is from two independent experiments, with the mean ratio ± SD of SUMOylated-versus total Ecm11 protein indicated.

Since the Zip4-Ecm11 interaction itself is not important for Zip4 ZMM function, we next assessed if it is involved in Ecm11 recruitment to chromatin. Indeed, ChIP-qPCR analyses revealed that Ecm11 was no longer recruited to all tested loci in both *zip4N919Q* and *ecm11LLDD* mutants (Fig. 3B and Supplemental Fig. S4C). This loss was further confirmed by Ecm11 and Red1 co-immunostaining of chromosome spreads, where *zip4N919Q* and *ecm11LLDD* cells showed no staining or discontinuous Ecm11 pattern, by contrast to wild type where 75% of meiotic cells showed continuous Ecm11 pattern (Fig. 3C-3D and Supplemental Fig. S5).

We next assessed the consequences of these Ecm11 loading defects on SC assembly, by Zip1 immunostaining of meiotic chromosome spreads. In wild-type cells at 5 h (pachytene stage), Zip1 staining was linear throughout the length of the chromosomes (Fig. 3E, upper panel). In contrast to wild type, but similar to *zip4*Δ and *ecm11*Δ, both *zip4N919Q and ecm11LLDD* mutants exhibited a discontinuous Zip1 pattern and decrease of the Zip1 fluorescence signal intensity (Fig. 3E-3F). In the interaction mutants, Zip1 localization defects were accompanied by the formation of Zip1 aggregates (polycomplexes), like in *ecm11*Δ (Fig. 3E, arrow -3G). Altogether, we showed that the Zip4-Ecm11 interaction is necessary for Ecm11 recruitment to chromosomes and normal SC assembly.

Previous studies have shown that Ecm11 is SUMOylated, depending on the Siz1 and Siz2 E3 ligases and that this is required for SC polymerization (Humphryes et al., 2013; Leung et al., 2015). However, we do not know if SUMOylation is linked to Ecm11 recruitment to chromosomes. Using our interaction mutant *zip4N919Q*, we found that Ecm11 SUMOylation levels were unchanged (Fig. 3H), clearly indicating that Ecm11 SUMOylation and its association to chromosomes concur independently to allow SC polymerization.

### Impaired Zip4-Ecm11 interaction increases homolog nondisjunction

Since Ecm11 and Gmc2 proteins were reported to influence to some extent DSB frequencies and CO distribution (Humphryes et al., 2013; Lee et al., 2021; Mu et al., 2020; Voelkel-Meiman et al., 2016), we investigated the function of the Zip4-Ecm11 interaction on recombination. We first measured CO frequency on two intervals on chromosome VIII (*CEN8-ARG4* and *ARG4-THR1*) by a fluorescent spore autonomous assay that also allows to measure homolog missegregation (MI nondisjunction) that can result from recombination defects (Thacker et al., 2011) (Fig. 4A). As expected for a *zmm* mutant, CO frequency in *zip4*Δ was decreased to about 28-35% of the wild type in the two intervals (Fig. 4B, and Supplemental Table S1). By contrast, *ecm11*Δ strain showed wild type CO levels in the *ARG4-THR1* interval and a slight but significant CO reduction (95% of wild type) in the *CEN8-ARG4* interval, confirming the interval-dependent effect of *ecm11*Δ. Similarly, the *zip4N919Q* interaction mutant showed wild type CO levels in the *ARG4-THR1* interval while it was reduced in the *CEN8-ARG4* interval, at an intermediate level between wild type and *zip4*Δ.

**Figure 4:**
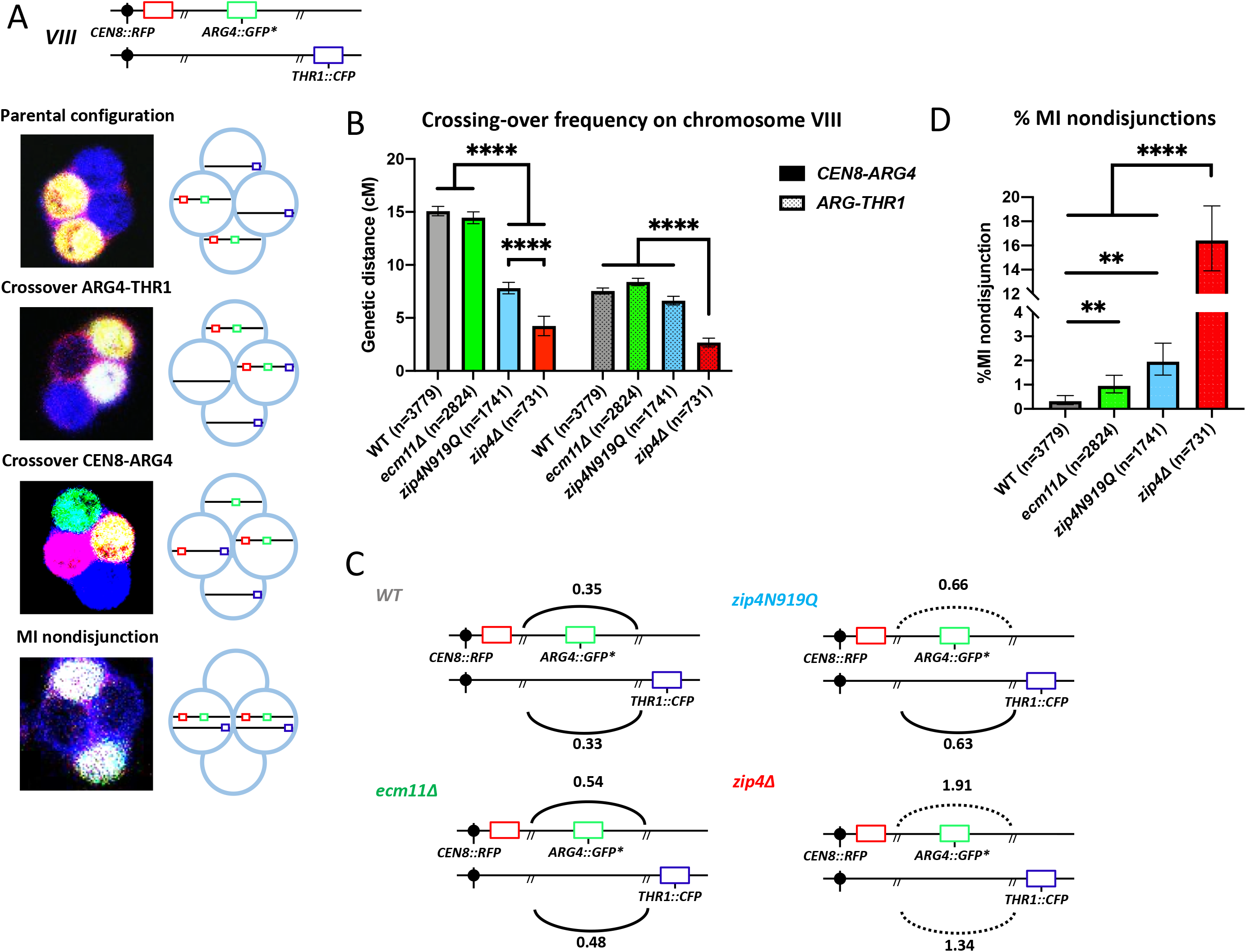
Effect of the different mutations on meiotic recombination and chromosome segregation. A. Illustration showing the location of the spore-autonomous reporters on chromosome VIII and the types of tetrads analyzed (Thacker et al., 2011). B. Crossing-over frequency measured in two genetic intervals *CEN8-ARG4* and *ARG4-THR1* on chromosome VIII. Genetic distances are plotted as cM ± SE for the indicated genotypes. **\*\*\***\*: p-value<0.0001, G-test. C. Interference between the two adjacent *CEN8-ARG4* and *ARG4-THR1* intervals calculated based on (Malkova et al., 2004) for the indicated genotypes. Solid line indicates that significant interference was observed. Dotted line indicates absence of significant interference. D. MI nondisjunction of chromosome VIII assessed by the spore-autonomous fluorescent reporter assay (see A). % MI nondisjunction ± 95 % CI is plotted. **: p-value<0.01, ****: p-value<0.0001, Fisher’s exact test.

We next assessed CO interference between the *CEN8-ARG4* and *ARG4-THR1* intervals (Fig. 4C). Interference was only slightly diminished in the *ecm11*Δ (0.51 vs 0.34 in wild-type), confirming previous studies (Lee et al., 2021; Voelkel-Meiman et al., 2016). Similarly, interference in the *zip4N919Q* mutant was slightly reduced (0.62), whereas it was completely abolished in *zip4*Δ (1.6) as expected for a *zmm* mutant (Fig. 4C). Therefore, the Zip4 mutant for interaction with Ecm11 behaves much more like a *ecm11*Δ mutant than a *zip4*Δ mutant, confirming the essential role of Ecm11 recruitment by Zip4 for Ecm11’s functions in SC assembly and recombination but not for the ZMM functions of Zip4.

Finally, using the spore fluorescent setup, we found that there was a low but significant increase of chromosome MI nondisjunction in both *ecm11*Δ (0.96 % ± 0.18 %) and *zip4N919Q* (1.95 % ± 0.33 %) compared to wildtype (0.32 % ± 0.09 %), which is much less than that seen in the *zip4*Δ mutant (16%) (Fig. 4D and Supplemental Table S1). This modest increase in nondisjunction may stem from the altered crossover frequency/distribution in the absence of Ecm11.

Overall, we conclude that impairing the interaction between Zip4 and Ecm11 mimics an *ecm11*Δ phenotype, confirming that Zip4 is responsible, in addition to its ZMM function, for all the functions of Ecm11 in SC assembly and recombination control.

### Artificially tethering interaction-deficient Ecm11 to Zip4 reinforces SC polymerization and accelerates meiotic progression

We next tested if artificially tethering the Ecm11LLDD mutant protein to Zip4 would be sufficient for SC polymerization and meiotic progression. For this, we fused Ecm11LLDD and Zip4 with FRB and FKPB12, respectively, to tether the two proteins upon rapamycin addition at 3.5 h in meiosis, just before the expected time of recombination (Fig. 5A). Interestingly, addition of rapamycin induced a faster meiotic progression, suggesting that facilitating Zip4-Ecm11 interaction may relax the checkpoint activated in the absence of the SC central element (Fig. 5B). We thus monitored SC polymerization by surface-spreading and Zip1 staining of meiotic cells and indeed, at all the time points tested, a strong increase of Zip1 fluorescence signal intensity was observed upon addition of rapamycin compared to the control condition (Fig. 5C-5D and Supplemental Fig. S6A-B). In addition, although many cells still contained Zip1 polycomplexes, their size was strongly decreased, consistent with better SC polymerization (Fig. 5C-5D and Supplemental Fig. S6C). We conclude that physically tethering Ecm11 to Zip4 is important for the incorporation of Zip1 within the SC and is able to partly compensate for the interaction defects of the Ecm11LLDD mutant. Therefore, our data suggest that rescuing Zip4-Ecm11 association facilitates polymerization of the transverse filament protein Zip1 and accelerates meiotic progression. Unexpectedly, tethering Zip4 to Ecm11 decreased spore viability and genetic distances, and increased homolog nondisjunction (Supplemental Fig. S6D-F). However, since meiotic progression was accelerated, tethering likely does not result in DSB repair defect, but most likely in decrease of DSB numbers, and therefore insufficient crossovers. We favor the hypothesis that unscheduled, early tethering of Zip4 to Ecm11 may trigger untimely, premature SC formation, and early inhibition of DSB formation, given the recently discovered function of SC polymerization to shut down DSBs (Mu et al., 2020).

**Figure 5:**
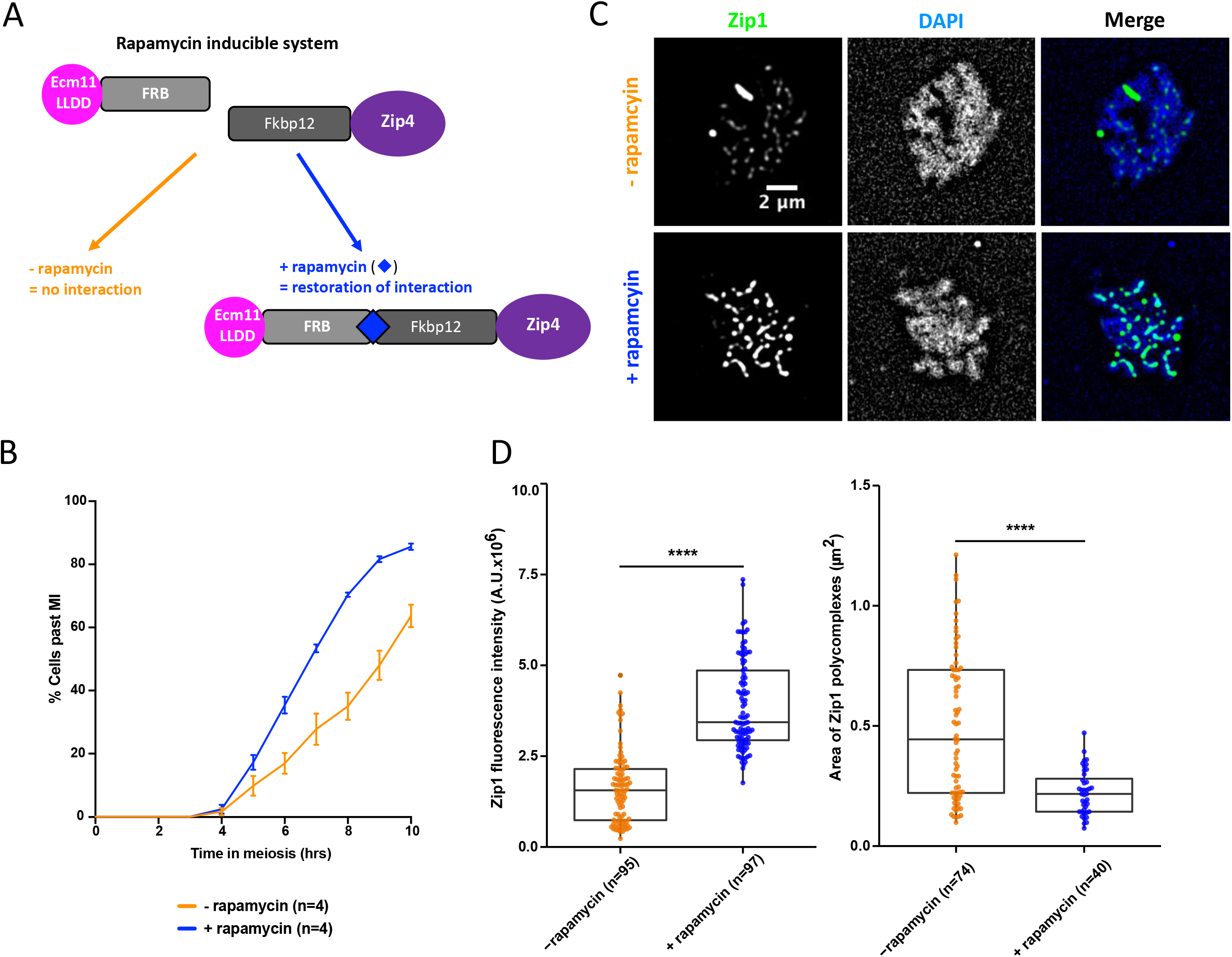
Forcing the interaction between Ecm11 and Zip4 is sufficient to restore both Ecm11 recruitment to chromosomes and synaptonemal complex assembly. A. Strategy to tether Ecm11LLDD fused to FRB domain to Zip4 fused to Fkpb12 domain by addition of rapamycin. B: Meiotic progression as assessed by DAPI staining of nuclei to monitor meiotic divisions. C. Zip1 localization on surface-spread chromosomes with (“+ rapamycin”) and without (“- rapamycin”) 1 µM rapamycin added at 3.5 h after meiotic induction. Pachytene stage nuclei are shown. Green: anti-Zip1; blue: DAPI. D. Left: quantification of Zip1 intensity observed in C. Right: quantification of polycomplexes area observed in C. Numbers of spreads are indicated for each condition. ****: p-value<0.0001, Wilcoxon test.

### The mouse Zip4 interacts with TEX12, a component of the SC central element, and Ecm11-Gmc2 show striking homology to TEX12-SYCE2

The whole ZZS complex (TEX11/Zip4-SHOC1/Zip2-SPO16) is present in mammals and is important for CO formation and fertility (Adelman and Petrini, 2008; Guiraldelli et al., 2018; Wang et al., 2001; Yang et al., 2008; Yatsenko et al., 2015; Yu et al., 2021; Zhang et al., 2018, 2019). Likewise, the SC overall structure is also conserved between budding yeast and mammals (Zickler and Kleckner, 2015). We therefore asked whether the interaction between Zip4 and the SC central element was conserved in mammals, by testing the interaction between mouse TEX11 and each of the five known proteins of the mouse SC central element: SYCE1, SYCE2, SYCE3, TEX12 and SIX6OS1 (Fraune et al., 2012; Gómez-H et al., 2016) (Fig. 6A). First, we recapitulated all the previously described interactions among the SC central element proteins by yeast two-hybrid, indicating that our constructs are functional for protein-protein interaction (Fig. 6A and Supplemental Table S2). The mouse TEX11 contains an aromatic-asparagine motif WN, as the yeast Zip4, in position 857-858 (Fig. 2C). In addition, a recent study in humans patients showed that the substitution of the Trp to Cys in this WN motif is associated with azoospermia (Sha et al., 2018). We thus generated a truncated TEX11 encompassing the C-terminal part of the protein (residues 637-947), named TEX11^Cter^, comprising the WN motif. Interestingly, we unveiled an interaction between TEX11^Cter^ and TEX12 (Fig. 6B), reminiscent of the Zip4-Ecm11 interaction in yeast. This suggests that the interaction between the ZMM protein Zip4/TEX11 and the central element of the SC may be conserved, and that TEX12 may be a functional homolog of Ecm11. The three-dimensional structures of human TEX12 and its close interacting partner, SYCE2, have been solved (PDB: 6R17) (Figure 6C) (Davies et al., 2012; Dunce et al., 2021). TEX12 is predicted to be SUMOylated on lysine 8, located at the very N-terminal extremity of the protein (see Materials and Methods), similarly to Ecm11 SUMOylation at lysine 5 (Humphryes et al., 2013). The similarity between TEX12 and Ecm11 is further strengthened by the coevolution patterns that are observed between Gmc2 and Ecm11 on one side and those between SYCE2 and TEX12 on the other side (Supplemental Fig. S7). Strikingly, although no evolutionary relationships could clearly connect the yeast and mammalian systems, their members are both predicted to interact through an anti-parallel followed by a parallel coiled-coil (Supplemental Fig. S7). This coevolution pattern is fully consistent with the structure of the SYCE2-TEX12 hetero-tetramer (Dunce et al., 2021) (Fig. 6C). Based on this experimental validation that the coevolution patterns for SYCE2-TEX12 are highly meaningful, we used the contacts predicted for the Ecm11-Gmc2 complex to generate a model of how the two proteins could interact with each other forming a tetrameric bundle likely to further self-assemble through regions flanking the canonical coiled-coil region (Supplemental Fig. S7 and Fig. 6D). Interestingly, the C and N-terminal extremities of TEX12 and SYCE2 appeared as essential for the complex to make fibers, consistent with a function for SC propagation (Fig. 6E). Pushing forward the analogy with TEX12-SYCE2 bundle, similar fibers may be formed by the Ecm11-Gmc2 complex through the conserved hydrophobic stretches upstream of the coiled-coil regions (Supplemental Fig. S7) and such structure could emanate from the SIC to catalyze SC polymerization by Zip1 (Fig. 6E).

**Figure 6:**
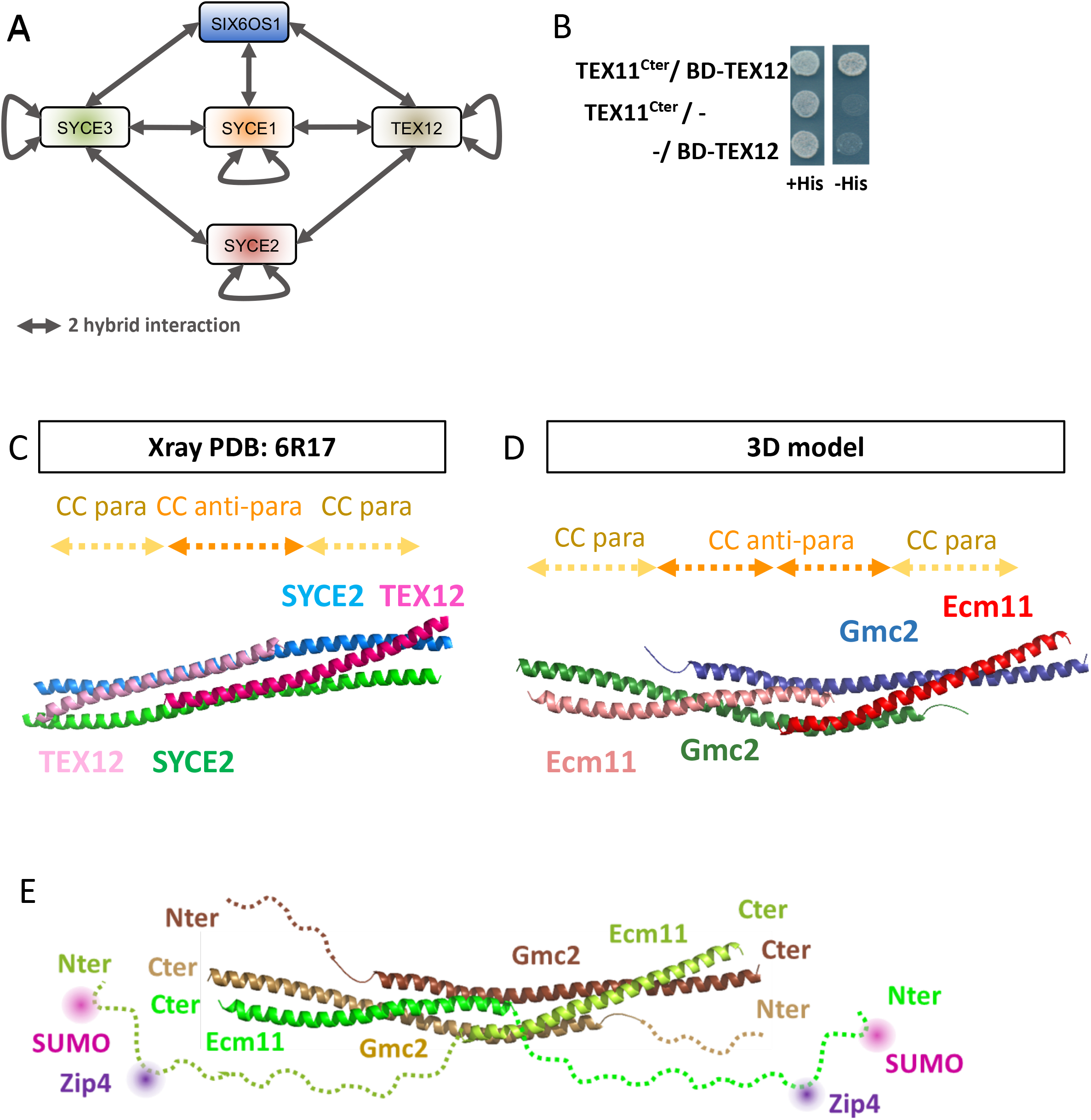
Mouse Zip4 (TEX11) interaction with the SC central element and analogies between yeast Ecm11-Gmc2 and mouse SYCE2-TEX12. A. Illustration showing the SC central element components in mouse and the two-hybrid interactions between them (see text). B. Yeast two-hybrid interaction analysis between mouse TEX11 and TEX12. C. Cartoon representation of the crystal structure of the SYCE2-TEX12 coiled-coils (PDB:6R17)(Dunce et al., 2021). SYCE2 is in blue and green and TEX12 in dark and light pink. The positions of the anti-parallel and parallel coiled-coil stretches are indicated by dashed arrows on top. D. 3D model of Ecm11-Gmc2. A model was built using Rosetta to fold the four subunits together under the co-evolution constraints. Gmc2 subunits are shown as blue and green cartoons while Ecm11 is shown as red and salmon cartoons. The locations of the parallel and anti-parallel stretches are indicated by dashed arrows on top (see also Supplemental Fig. S7). E. Similar model as in 6D integrating the Nter regions of Ecm11 and Gmc2, highlighting the SUMOylation (pink circle) and Zip4 (dark purple circle) interaction sites of Ecm11.

## Discussion

Several studies point to a close relationship between crossover sites and sites of SC nucleation (Pyatnitskaya et al., 2019). However, the connection between these two important processes remained elusive. Here, we described a direct and functional interaction between the ZMM protein Zip4 and Ecm11, a component of the SC central element, providing the physical link between crossovers and SC polymerization.

### Zip4 is an interface protein that integrates signals from both crossover and synapsis promoting factors

Zip4 is a protein with repetitive TPR domains, motifs that are common in scaffold proteins and exhibit a wide range of molecular recognition modes (D’Andrea and Regan, 2003; Perez-Riba and Itzhaki, 2019). An interesting property of some TPR proteins is their ability to orchestrate different activities by integrating signals from multiple interacting partners. Several pieces of evidence point out to such a role for the Zip4 protein. Firstly, on the “ZMM side”, Zip4 interacts directly with its ZMM partners Zip2 and Spo16 to form the ZZS complex. Within this complex, a domain of Zip2 forms with Spo16 an XPF-ERCC1-like module that recognizes DNA joint molecules (Arora and Corbett, 2019; De Muyt et al., 2018). The role of Zip4 in this complex is not well understood but Zip4 is important for Zip2 stability, and may act as a chaperone for Zip2 and Spo16, reinforcing their DNA recognition activity (De Muyt et al., 2018). Secondly, the other ZMM proteins, SUMO/Ubiquitin ligase Zip3 and MutS*γ* have also been reported to colocalize and interact with Zip4, suggesting that Zip4 integrates multiple ZMM activities to consolidate joint molecules intermediates and promote CO formation (De Muyt et al., 2018; Shinohara et al., 2008).

In addition to ZMMs, Zip4 interacts with components of the SC. In budding yeast, a connection between Zip4 and the synaptonemal complex was first identified via a direct interaction with Red1, the axial element of the SC (De Muyt et al., 2018). This seems conserved in mammals since the Zip4 ortholog, TEX11, interacts with the SC axial element SYCP2 (Yang et al., 2008). We showed here that Zip4 also binds to the SC central element Ecm11 and Gmc2 proteins, suggesting that Zip4 is tightly connected to SC proteins through multiple interactions. Interestingly, the axial and central elements of the SC are separated from each other by 50 nm, suggesting that Zip4 is present in two different locations within the SC. Based on what is known about the temporal dynamics of recombination intermediates during the successive steps of recombination, we envision that Zip4, bound on recombination intermediates through the Zip2-Spo16 module, may first interact with the axial element (via Red1), at an early recombination step, and would then be translocated to the future central element location, between the axes, at a later step of recombination, to seed SC nucleation *via* its interaction with Ecm11. Such dynamics would result in bringing “miniature axes” (or bridges), containing Red1, from parental chromosomes into the inter-axis region, as proposed in *Sordaria* (Dubois et al., 2019).

Several lines of evidence suggest that the SC emerges from ZMM-bound sites. In budding yeast, *Sordaria* and mouse, SC initiation sites often colocalize with ZMM proteins (Agarwal and Roeder, 2000; Dubois et al., 2019; Reynolds et al., 2013; Tsubouchi et al., 2006) and decrease in number in mutants with reduced DSB numbers, while synapsis defects are increased (Henderson and Keeney, 2004; Kauppi et al., 2013; Tessé et al., 2003, 2017). This suggests that a minimum number of ZMM/SC nucleation sites are required for full homolog synapsis (Tsubouchi et al., 2006). Since the SC transverse filament protein Zip1 is also a ZMM protein, it was an obvious candidate for the initial recruitment of Ecm11. Moreover, an N-terminal deletion mutant *zip1N1* has a similar phenotype to *ecm11*Δ and Ecm11-Gmc2 colocalize with Zip1 during synapsis initiation and completion (Humphryes et al., 2013; Tung and Roeder, 1998; Voelkel-Meiman et al., 2016). However, we found no evidence of interaction between Zip1 and Ecm11 or Gmc2 in our Y2H experiments. In addition, Ecm11 foci are still visible in a *zip1*Δ mutant (Humphryes et al., 2013) and Ecm11 still associates to DSB hotspots in our ChIP experiments, implying that Zip1 is not required for the initial SC assembly from the ZMM nucleation sites. Instead, we provide a body of evidence that Zip4, through its direct interaction with Ecm11, plays a pivotal role promoting synapsis at these ZMM binding sites: (i) Ecm11 shows a pattern similar to that of ZMMs, binding both DSB and axis sites; (ii) Ecm11 localization at DSB sites strictly requires Zip4 protein, in agreement with the absence of Ecm11 foci in *zip4*Δ mutant, but not in other tested *zmm* mutants (Humphryes et al., 2013); (iii) mutations altering the interaction between Zip4 and Ecm11, *zip4N919Q and ecm11LLDD*, result in defective SC assembly and in polycomplex formation, in a manner akin to *zip4Δ and ecm11*Δ; (iv) tethering Zip4 and a mutated interaction-defective Ecm11 is sufficient to restore SC assembly and faster meiotic progression. In budding yeast, Ecm11 acts in complex with Gmc2 during SC polymerization (Humphryes et al., 2013). Interestingly, the Ecm11LLDD mutated protein keeps its ability to form a heterodimer with Gmc2. Moreover, Gmc2 also interacts with Zip4 in yeast two-hybrid, suggesting that Zip4 may promote SC assembly by depositing a pre-formed Ecm11-Gmc2 complex. Finally, we can envision that Zip4 coordinates signals at the same time through simultaneous interactions between different TPR motifs present throughout its length and its proteins partners (including Zip2, Zip3, Msh5, Red1, Ecm11 and Gmc2). It will be of interest to identify the role of all the sites docking Zip4 to its described partners.

### Spatio-temporal coupling of crossovers and SC assembly

Given the importance of Zip4 in the recognition of DNA joint molecules through the ZZS module and in SC assembly via Ecm11 (and Gmc2) interaction, and to integrate all present and past results, we propose the following model for Zip4 mechanism of action (Fig. 7): 1) After DSB formation, the ZZS complex associates with recombination intermediates via the XPF-ERCC1-like DNA recognition module (De Muyt et al., 2018), and with the axis component Red1. Other ZMMs, including Zip1, also bind recombination intermediates. 2) Then, still bound on recombination intermediates, the ZZS complex transits from the axis region towards the inter-axis region, leading to the formation of chromosomal bridges that progressively align the parental chromosomes (De Muyt et al., 2018; Dubois et al., 2019; Pyatnitskaya et al., 2019). In the meantime, Zip4 helps to bring Ecm11-Gmc2 at these sites by direct protein-protein interaction. 3) The Ecm11-Gmc2 complex helps initiate the polymerization of the surrounding Zip1. It is at this time that a “synapsis initiation complex” is created and the SC will start to emanate from this nucleation zone, through Zip1 polymerization. 4) Finally, as suggested recently, this SC polymerization exerts a negative feedback on *de novo* DSBs formation, and therefore locally affects crossover frequencies (Lee et al., 2021; Mu et al., 2020; Thacker et al., 2014; Voelkel-Meiman et al., 2016) (Fig. 7). This mechanism of regulation starting from crossover-designated sites would be an elegant way for the cell to fine-tune CO patterning by shutting down DSBs locally through the propagation of the SC along chromosomes.

**Figure 7:**
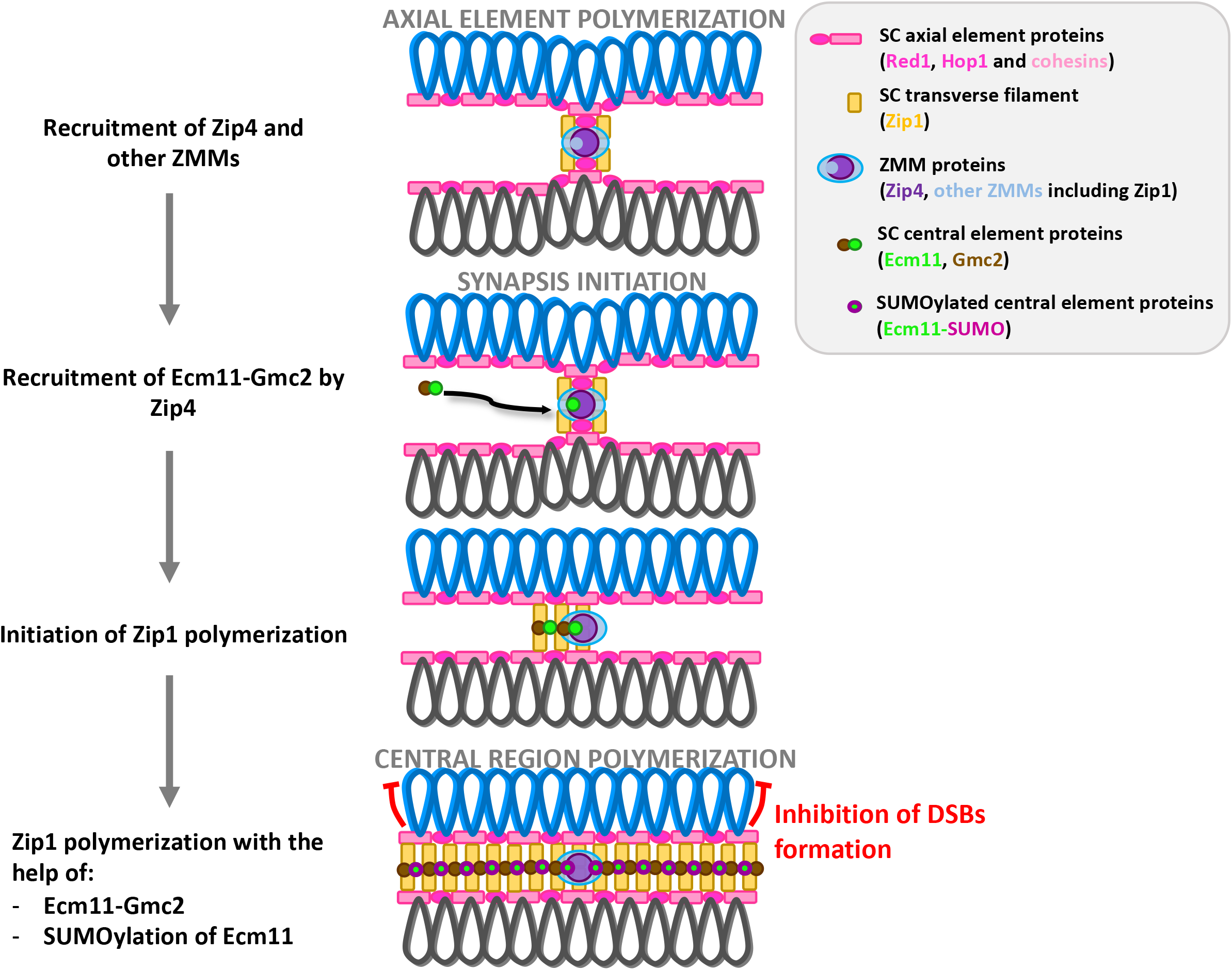
Model for the link between crossover sites and SC assembly. The model is based on our study of Zip4-Ecm11 interaction and published studies (see text). First, axial element polymerizes and SICs are formed after the transition of ZMMs (including Zip4, in dark purple) to the inter-axis region. The Ecm11 (green)-Gmc2 (brown) heterodimer is brought to the SIC through its interaction with Zip4, which initiates the polymerization of the TF Zip1 (purple). Polymerization of the central region composed of the TF Zip1 and the central element Ecm11-Gmc2 progresses, closely aligning the homologs at a 100 nm distance. PolySUMOylation of Ecm11 (indicated by pink circle) triggered by the TF assembly exerts a positive feedback on the central region polymerization (Leung et al., 2015). SC central region assembly inhibits the formation of de-novo DSBs, thus avoiding additional break and repair in already synapsed regions.

### The relationship between crossovers and SC assembly in other species

Like in budding yeast, in mice and plants, the absence of DSB or efficient interhomolog repair processes leads to synapsis defects suggesting that synapsis initiation depends on the total number of interhomolog interactions (Cahoon and Hawley, 2016; Mercier et al., 2015; Pyatnitskaya et al., 2019). It is currently unknown whether a protein complex similar to the SIC is required for the initiation of SC polymerization at these sites of interhomolog engagement. However, since mouse *zmm* mutants show synapsis defects, ZMM proteins could participate in the initiation of SC formation, although the different extent of synapsis defects observed among zmm mutants suggests that the absence of some ZMM might be concealed by a second mechanism based on homology-independent SC extension, known as synapsis adjustment (Zickler and Kleckner, 1999). Finally, contrary to budding yeast, whereas ZMM proteins are still detected between homolog axes, the SC central element proteins SYCE1/2/3 and TEX12 are no longer detected on chromosomes in *Sycp1^-/-^* (Hamer et al., 2006; Schramm et al., 2011). The central element proteins may have a different mode of recruitment and/or their abundance is too low to be detected by conventional microscopy, if they form only dots, as in budding yeast.

In plants, SC polymerization seems less dependent on the CO-mediated interhomolog engagement, since *zmm* mutants does not have apparent synapsis defects (Mercier et al., 2015), but this does not mean that SC polymerization does not initiate from ZMM-bound sites in wild type. In addition, kinetics of SC assembly and synergistic effects of ZMM mutations have not been thoroughly tested. Indeed, combination of both *zip4* and *mer3* mutations leads to severe synapsis defects in rice, suggesting that ZMM proteins might have redundant roles for SC loading in plants (Shen et al., 2012).

In contrast to budding yeast, plants and mammals, some species use a recombination-independent mode of initiating SC polymerization. In particular, in the worm *C. elegans*, it starts from telomeres and in the fly *Drosophila*, it starts from centromeres (Christophorou et al., 2013; Dernburg et al., 1998; MacQueen et al., 2005; McKim et al., 1998). Interestingly, these species lack many of the ZMM proteins including Zip4, Zip2 and Spo16, maybe resulting from the absence of selective pressure for CO-designated interhomolog engagement for SC initiation.

### Concluding remarks

Recent studies in yeast and plants showed the importance of close homolog juxtaposition by the SC to control recombination frequency and crossover distribution (Capilla-Pérez et al., 2021; France et al., 2021; Lee et al., 2021; Mu et al., 2020). We propose that this control is initiated by the direct interaction between Zip4 and Ecm11. It will be important to understand the interplay between this coupling mechanism and the mechanism of the initial deposition of Zip1, which requires Mek1 phosphorylation, to coordinate SC assembly (Chen et al., 2015). Finally, further investigations on the relationship between the ZMM-dependent CO formation and the SC dynamics in different model organisms will be needed to uncover both their conserved as well as distinct features and reveal how it could impact human fertility, given the involvement of TEX11 mutations in patients with azoospermia.

## STAR Methods

### Yeast manipulation

All yeast strains are derivatives of the *SK1* background except those used for two-hybrid experiments and for ChIP-seq spike-in control. Their complete genotype and their use in different figures are in Supplemental Table S3. All experiments were performed at 30 °C. For synchronous meiosis, cells were grown in SPS presporulation medium and transferred to 1% potassium acetate with vigorous shaking at 30 as described (Murakami et al., 2009). For all strains, spore viability was measured after sporulation on solid sporulation medium for two days at 30 C.

### Yeast strains construction

Yeast strains were obtained by direct transformation or crossing to obtain the desired genotype. Site directed mutagenesis and C-terminal deletions were introduced by PCR. All transformants were confirmed using PCR discriminating between correct and incorrect integrations and sequencing for epitope tag insertion or mutagenesis. The functionality of the tagged proteins was measured by spore viability assays. All tagged proteins were functional.

### Sequence analyses and modelling of Zip4, Ecm11 and Gmc2 structures

Full-length homologous sequences of Zip4, Ecm11, Gmc2, TEX12 and SYCE2 were retrieved using PSI-BLAST iterations on the nr database, gathering 862, 916, 824, 165 and 184 sequences, respectively. Multiple sequence alignments were generated for these sets of sequences using MAFFT (Katoh and Standley, 2013) and represented using Jalview (Waterhouse et al., 2009). Co-MSA for the Gmc2-Ecm11 and SYCE2-TEX12 were obtained by selecting a single sequence per species selecting the hit of lowest e-value and by concatenating the alignments resulting in a co-MSA of 451 and 135 sequences, respectively. These alignments were used as input of the RaptorX contact prediction (Wang et al., 2017) to predict the contact maps within and between the pairs of proteins. A 3D model of Zip4 was generated using the latest version of the RoseTTAFold server combining coevolution and deep learning approaches for the prediction of 3D monomeric structures (Baek et al., 2021). Analyses of the SUMOylation sites were performed using the Jassa server (Beauclair et al., 2015) and those of the coiled-coils were performed using PCOILS as implemented in the MPI Bioinformatics Toolkit server (Lupas et al., 1991).

### Yeast two-hybrid analyses

Strains expressing *ZMMs* are described in (De Muyt et al., 2018). *ECM11* and *GMC2* were PCR-amplified from SK1 genomic DNA. Site-directed mutations were introduced by fusion of PCR products. Full-length mouse *Tex11*, *Tex12*, *Syce1*, *Syce2*, *Syce3*, *Six6os1* were PCR-amplified from mouse testis cDNA, a gift from D. Bourc’his. PCR products were cloned in plasmids derived from the 2 hybrid vectors pGADT7 or pGADCg (GAL4-activating domain) and pGBKT7 or pGBKCg (GAL4-binding domain), creating N- or C-terminal fusions and transformed in yeast haploid strains Y187 and AH109 (Clontech), respectively. Yeast two-hybrid assays were performed and interactions scored on selective media exactly as described in (Duroc et al., 2017).

### Analysis of crossover frequencies

Diploids were sporulated in liquid medium, and recombination between fluorescent markers on chromosome VIII was scored after 24 h sporulation, by microscopy analysis, as described previously (Thacker et al., 2011). Two independent sets of each strain were combined and at least 730 tetrads were scored for crossovers in two test intervals and for MI-nondisjunction events. Genetic distances in the *CEN8-ARG4* and *ARG4-THR1* intervals were calculated from the distribution of parental ditype (PD), nonparental ditype (NPD), and tetratype (T) tetrads and genetic distances (cM) were calculated using the Perkins equation: cM=(100 (6NPD + T))/(2(PD + NPD + T)). SEs of genetic distances were calculated using Stahl Lab Online Tools https://elizabethhousworth.com/StahlLabOnlineTools/.

### Cytology

For cytology, 1×10^8^ cells were harvested at the indicated time-point and yeast chromosome spreads were prepared as described in (Grubb et al. 2015). Primary antibodies used were mouse monoclonal 9E11 anti-myc antibody (dilution 1:200), rabbit polyclonal anti-Zip1 antibody (sc-33733, SantaCruz Biotech, dilution 1:100) and rabbit monoclonal anti-Red1 antibody (#16441, Gift from N. Hollingsworth, dilution 1:200). The secondary antibodies were Alexa488-conjugated goat anti-rabbit (A-11008, Thermo Fischer Scientific; dilution 1:200), Alexa568-conjugated goat anti-mouse (A-11004, Thermo Fischer Scientific; dilution 1:200). Chromosomal DNA was stained by 4,6-diamidino-2-phenylindole (DAPI). Fluorescence images were visualized and acquired using the Deltavision IX70 system (Applied Precision), objective 100X and softWoRx imaging software. Images were processed by deconvolution using the constrained iterative deconvolution algorithm within softWoRx. Image analysis and signal quantification was performed using the Fiji software and R-scripts. Fluorescence intensity was measured as the sum of pixel density of Zip1 stretches.

### TCA extraction and Western blot analysis

Protein extracts were prepared by trichloroacetic acid (TCA) precipitation method. 1.5 mL of sporulating cell culture was harvested and pellet was immediately frozen in liquid nitrogen. Cells were resuspended in 100 μL of ice-cold NaOH solution (1.85 N NaOH, 7.5% β-mercaptoethanol) and incubated for 10 min on ice. Samples were then mixed with 30 μL of ice-cold TCA 50% and incubated for 10 min on ice. Cell suspension was then harvested for 5 min at 15000 g at 4°C and the pellet was resuspended in 100 μL of loading buffer (55 mM Tris pH 6.8, 6.6 M Urea, 4.2% SDS, 0.083 mM EDTA, 0.001% bromophenol blue, 1.5% β-mercaptoethanol). Protein samples were dipped in liquid nitrogen and then incubated at 65°C for 3 min. Samples were centrifuged 5 min at 20000 g and the supernatant was kept at -80°C. Samples were loaded on precast acrylamide gel (4-12% Bis-Tris gel (Invitrogen)) and transferred on PVDF membrane in MOPS SDS Running Buffer (Life Technologies). Proteins were detected using mouse monoclonal M2 anti-Flag (F1804, Sigma, dilution 1:1000), mouse monoclonal 9E11 anti-myc (dilution 1:500) or rabbit monoclonal anti-TAP antibody (CAB1001, Invitrogen, dilution 1:2000). For normalization, mouse monoclonal anti-Pgk1 antibody was used (459250, Invitrogen, 1:3000). Image acquisition was performed with Chemidoc system (Biorad). To quantify protein levels, the band intensity in each lane was measured by the ImageLab software and divided by the corresponding Pgk1 band intensity in the same lane.

### Co-immunoprecipitation

1.2×10^9^ cells were harvested and 1mM of PMSF was added. Cells were washed once with PBS, and lyzed in 3 ml lysis buffer (20 mM HEPES/KOH pH7.5; 150 mM NaCl; 0.5% Triton X-100; 10% Glycerol; 1 mM MgCl2; 2 mM EDTA; 1 mM PMSF; 1X Complete Mini EDTA-Free (Roche); 1X PhosSTOP (Roche) with 0.5 mm zirconium/silica beads (Biospec Products, Bartlesville, OK) three times for 30s in a Fastprep instrument (MP Biomedicals, Santa Ana, CA). The lysate was incubated 1 h at 4°C with 125 U/mL of benzonase. 100 μL of PanMouse IgG magnetic beads (Thermo Scientific) were washed 1:1 with lysis buffer, preincubated in 100 μg/mL BSA in lysis buffer for 2 h at 4°C and then washed twice with 1:1 lysis buffer. The lysate was cleared by centrifugation at 13,000 g for 5 min and incubated overnight at 4°C with washed PanMouse IgG magnetic beads. The magnetic beads were washed four times with 1 mL of wash buffer (20 mM HEPES/KOH pH7.5; 150 mM NaCl; 0.5% Triton X-100; 5% Glycerol; 1 mM MgCl2; 2 mM EDTA; 1 mM PMSF; 1X Complete Mini EDTA-Free (Roche); 1X Phos-STOP (Roche)). The beads were resuspended in 30 μL of TEV-C buffer (20 mM Tris/HCl pH 8; 0.5 mM EDTA; 150 mM NaCl; 0.1% NP-40; 5% glycerol; 1 mM MgCl2; 1 mM DTT) with 4 μL TEV protease (1 mg/mL) and incubated for 2 h at 23°C under agitation. The eluate was transferred to a new tube. After washing, beads were resuspended in 25 μl of 2x SDS protein sample buffer. Beads eluate was heated at 95°C for 3 min and loaded on acrylamide gel (4-12% Bis-Tris gel (Invitrogen)) and run in MOPS SDS Running Buffer (Life Technologies). Proteins were then transferred to PVDF membrane using Trans-Blot® Turbo™ Transfer System (Biorad) at 2.5 A constant, up to 25 V for 10 min. Proteins were detected using mouse monoclonal M2 anti-Flag (F1804, Sigma, dilution 1:1000) or rabbit monoclonal anti-TAP antibody (CAB1001, Invitrogen, dilution 1:2000). Signal was detected using the SuperSignal West Pico or Femto Chemiluminescent Substrate (ThermoFisher). Images were acquired using Chemidoc system (Biorad). Signal was analyzed with ImageLab software. Results were presented as % INPUT band after subtracting the untagged strain signal and normalizing by TAP-tagged protein level.

### Chromatin immunoprecipitation

For each meiotic time point, 2×10^8^ cells were processed as described in (Duroc et al., 2017) except that before use, magnetic beads were blocked with 5 μg/μL BSA for 4 h at 4°C. Quantitative PCR was performed from the immunoprecipitated DNA or the whole-cell extract using a QuantStudio 5 (Applied Biosystems, Thermo Scientific) and analysed as described (Duroc et al., 2017). Results were expressed as % of DNA in the total input present in the immunoprecipitated sample. Primers for *GAT1*, *BUD23*, *HIS4LEU2*, *ERG1*, *AXIS* and *NFT1* loci have been described (Sanchez et al., 2020). For ChIP-seq experiments, 1×10^9^ cells were processed as described (De Muyt et al., 2018; Sanchez and Borde, 2021; Sanchez et al., 2020) except that, for spike-in normalization, 1×10^8^ (10%) *S. mikatae* cells of a single meiotic culture, harvested at 4 h in meiosis and fixed using the same procedure as for *S. cerevisiae*, were added to each sample before processing.

### Illumina sequencing of ChIP DNA and read normalisation

Purified DNA was sequenced using an Illumina NovaSeq 6000 instrument following the Illumina TruSeq procedure, generating paired-end 100 base-pair (bp) reads for Ecm11 in wild-type, *zip1*Δ, *zip4*Δ and untagged anti-Flag ChIP. Each experiment was performed in two independent replicates. Reads were aligned to the SaccCer2 *S. cerevisiae* S288C genome exactly as described (Sanchez et al., 2020), and to the *S. mikatae* genome assembly (Scannell et al., 2011). Reads that aligned on the *S. cerevisiae* genome but not on *S. mikatae* were defined as the experimental reads. For defining the spike-in normalization factor, we then determined the number of reads that did not align on the *S.cerevisiae* genome but aligned to the *S. mikatae* genome assembly (Scannell et al., 2011), generating the spike-in reads. The aligned experimental reads from independent replicates were then combined using MergeSamFiles to generate a single Bam file. Next, each Bam file was converted to bigwig format using deepTools bamCoverage, with a binsize of 1, a smoothing window of 200 bp and a normalization factor “2”, obtained as follows: for each sample, the number of experimental reads was first divided by the number of spike-in reads, giving scaling factor “1”. Then, the factor 1 of each sample was divided by the mean untagged sample coverage (290), giving scaling factor 2. Finally, for each bigwig file obtained, the scaled untagged sample was subtracted from the scaled tagged sample. These values were used for Fig. 1C,D. Sequencing data were deposited at the NCBI Gene Expression Omnibus database with the accession numbers GSE177033. Peaks for Red1 ChIP-seq and Spo11 oligonucleotides were from (Sun et al., 2015; Zhu and Keeney, 2015), respectively.

## Acknowledgments

We thank N. Hollingsworth for the anti-Red1 antibody, A. Hochwagen for the SK1 *fpr1 tor1-1* mutant strains, D. Bourc’his for the mouse testis cDNA, and Institut Curie NGS platform, supported by grants from ANR-10-EQPX-03, ANR10-INBS-09-08 and from the Cancéropôle Ile-de-France. This work was supported by Institut Curie and CNRS, by Agence Nationale de la Recherche (ANR-15-CE11-0011), Fondation ARC (PJA20171206487), Ligue Contre le Cancer (5FI13573TAZP). A. P. was a recipient of a predoctoral funding from the Fondation pour la Recherche Médicale.

## Author contributions

A.D.M. and V.B. supervised the study. A.P. and A.D.M. performed the experiments. J.A. and R.G. designed all the protein-protein interaction mutants and performed the structure predictions. A.P., A.D.M. and V.B. wrote the paper, with input from all the authors.

## Supplemental Tables

**Table S1:** Fluorescent spore assay data (Xcel spreadsheet).

**Table S2:** Yeast two-hybrid interactions between mammalian SC central element proteins and TEX11 (Xcel spreadsheet).

**Table S3:**
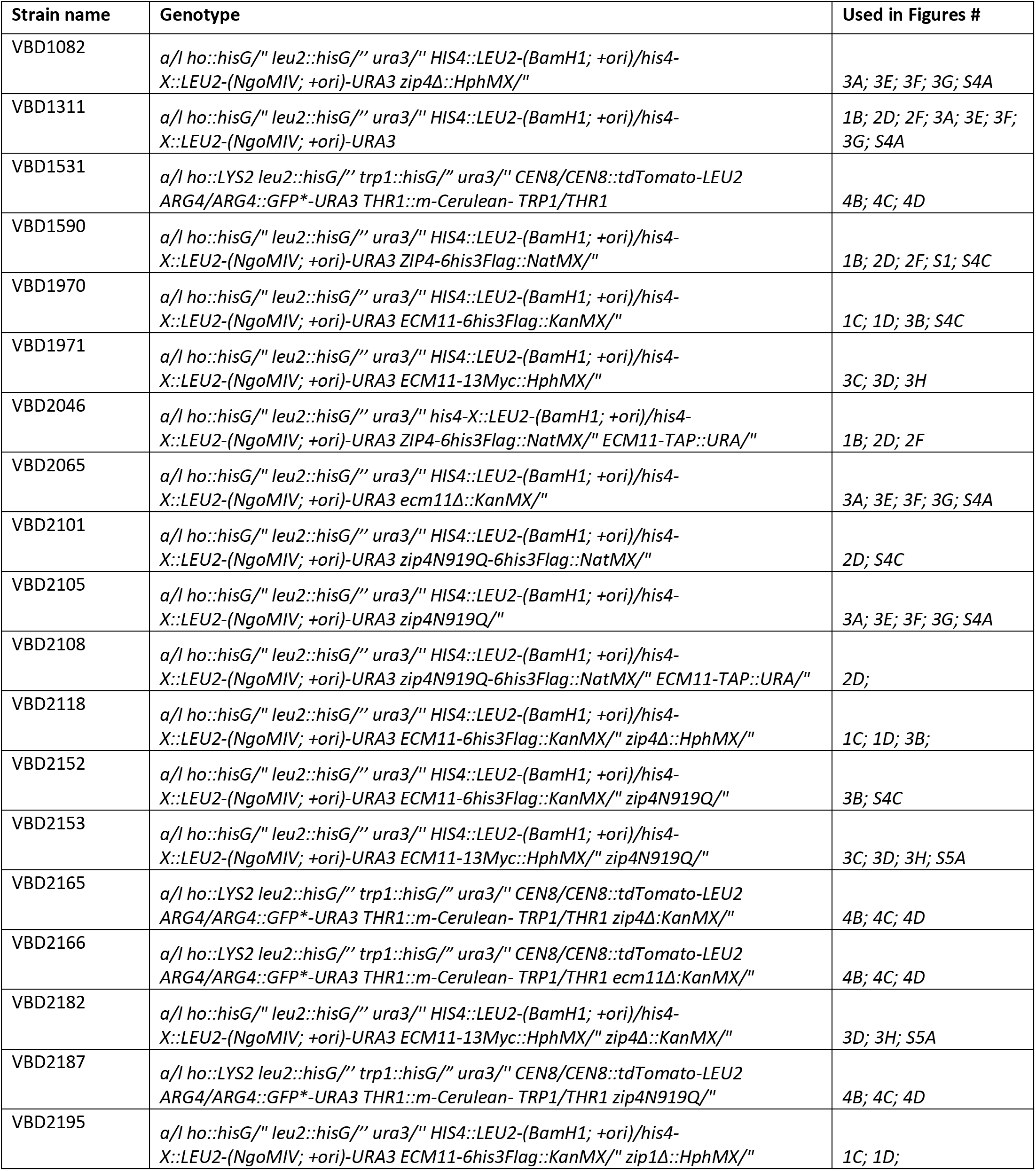
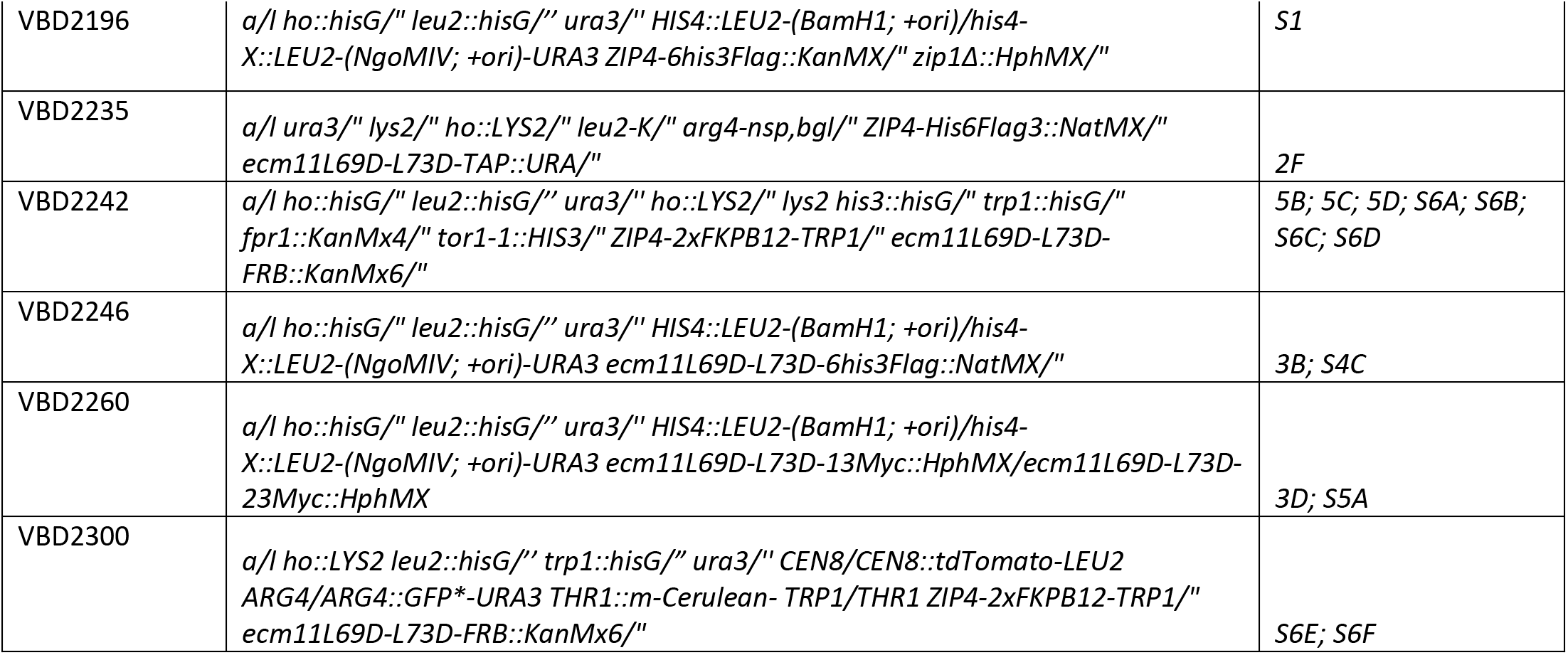
Yeast strains used.

## Supplemental figures legends

**Figure S1:**
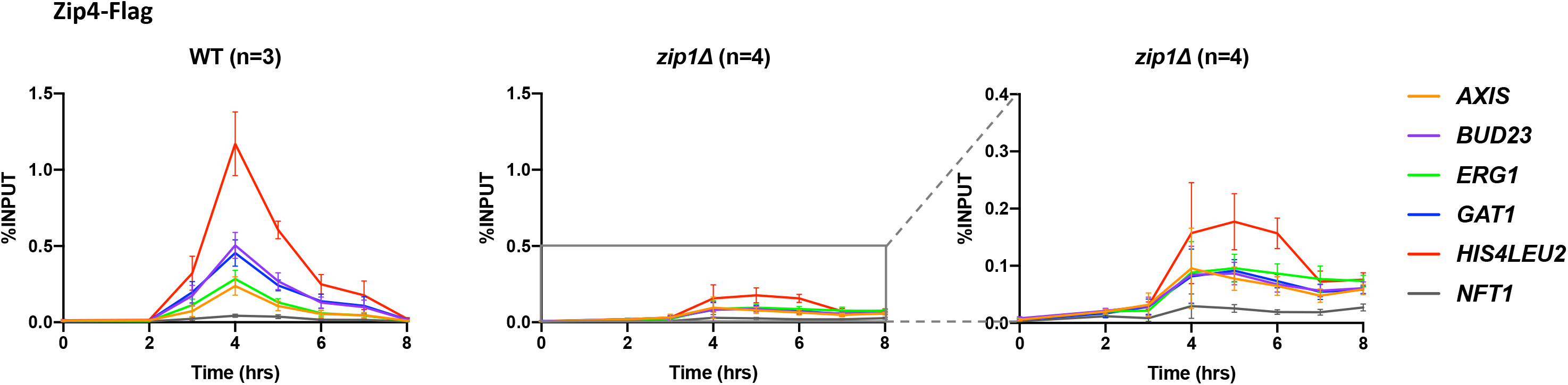
Zip4 binding to chromosomes is reduced in absence of Zip1 protein. ChIP monitoring of Zip4-Flag association with different chromosomal regions, measured by qPCR using primers that cover the indicated regions. Values are the mean ± SEM of at least three independent experiments. The graph on the right represents a magnification of the central graph.

**Figure S2:**
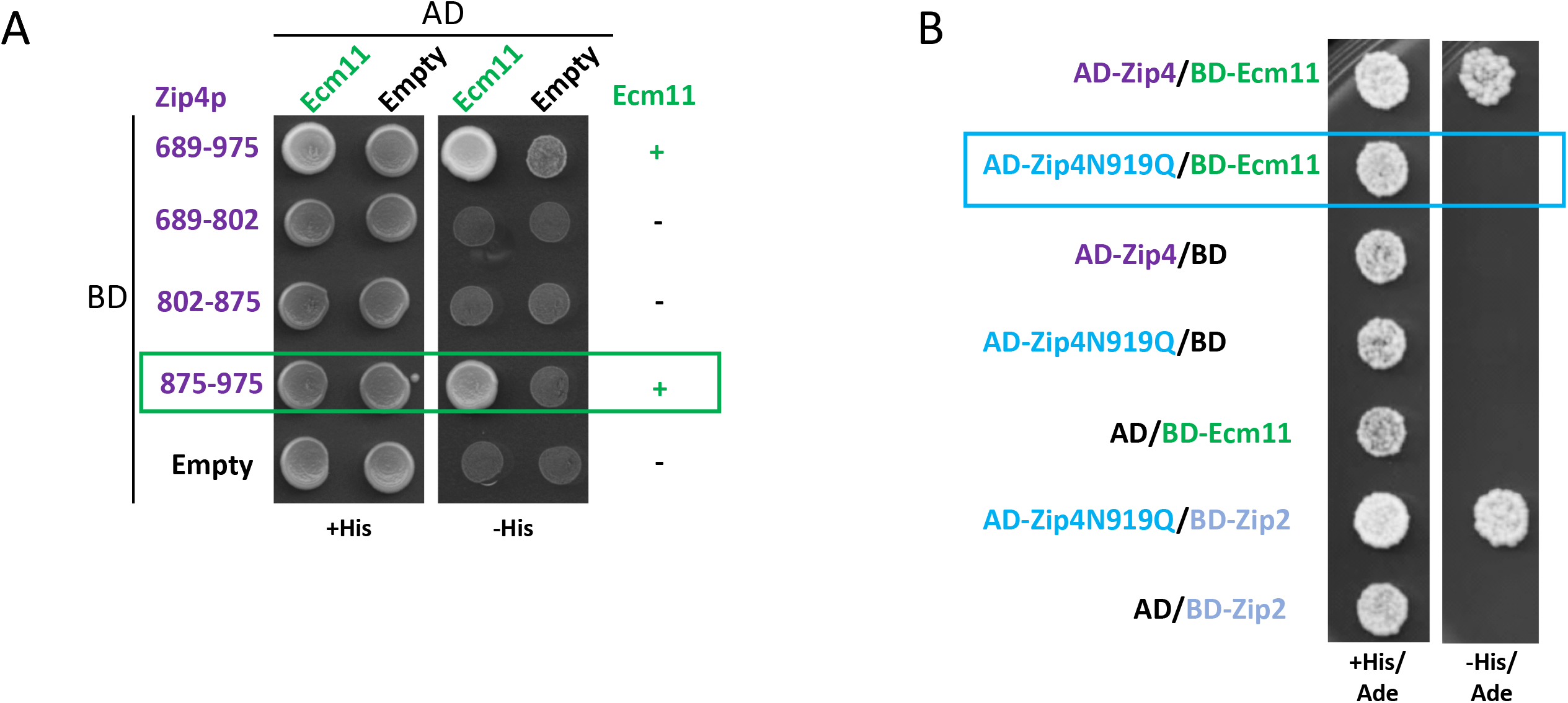
Zip4 interacts with Ecm11 through an aromatic-asparagine motif. A. Yeast two-hybrid interaction analysis between truncated Zip4 and Ecm11. Preys and baits are fused with the GAL4 Activation Domain (GAL4-AD) and with the GAL4 DNA-Binding Domain (GAL4-BD), respectively. The green frame indicates the interaction between Zip4-875-975 and Ecm11. B. Yeast two-hybrid interaction analysis between Zip4 and Ecm11/Zip2. The blue frame indicates the absence of interaction between Zip4N919Q and Ecm11. A-B: The interaction is revealed by growth on the selective –His/Ade medium.

**Figure S3:**
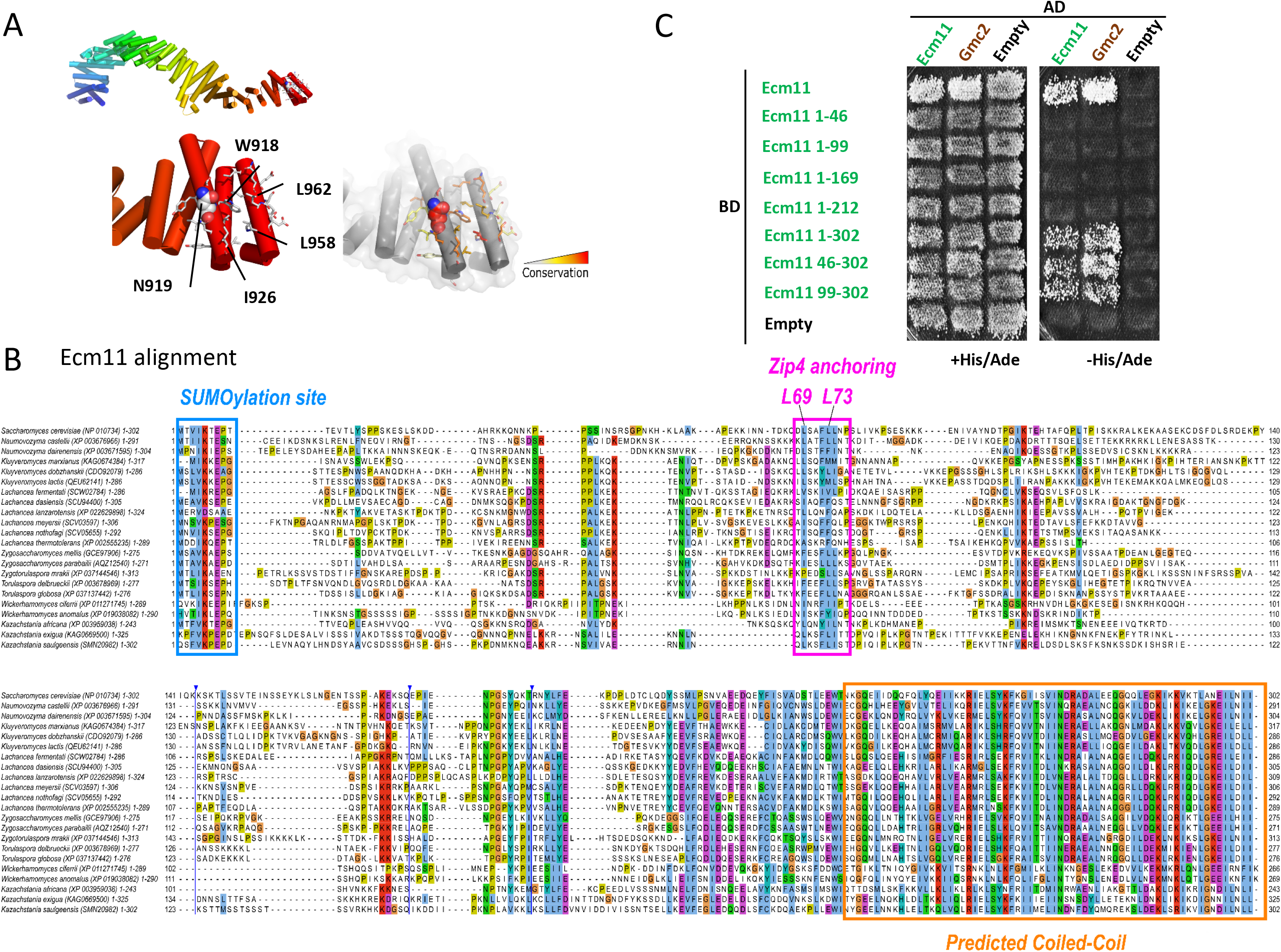
Delineation of the Ecm11 region interacting with Zip4. A. Illustration of the predicted structure of Zip4 WN motif interacting with Ecm11. B. Multiple sequence alignment gathering homologs of Ecm11 in budding yeasts of the *Saccharomycetaceae* family (22 sequences with their NCBI identifiers and delimitation index indicated). The conserved SUMOylation site containing the modified Lysine 5 is highlighted in the cyan box. The N-terminal region interacting with Zip4 and predicted to adopt a small helical conformation is boxed in magenta with the conserved positions of L69 and L73 highlighted. The C-terminal region predicted to form a coiled-coil over 63 residues is indicated by the orange box. Blue vertical lines indicate the positions of long insertions present in only a few homologs of *S. cerevisiae* Ecm11 which were masked for the sake of compact representation. C. Yeast two-hybrid self-interaction analysis of Ecm11. Same legend as in Fig. S2

**Figure S4:**
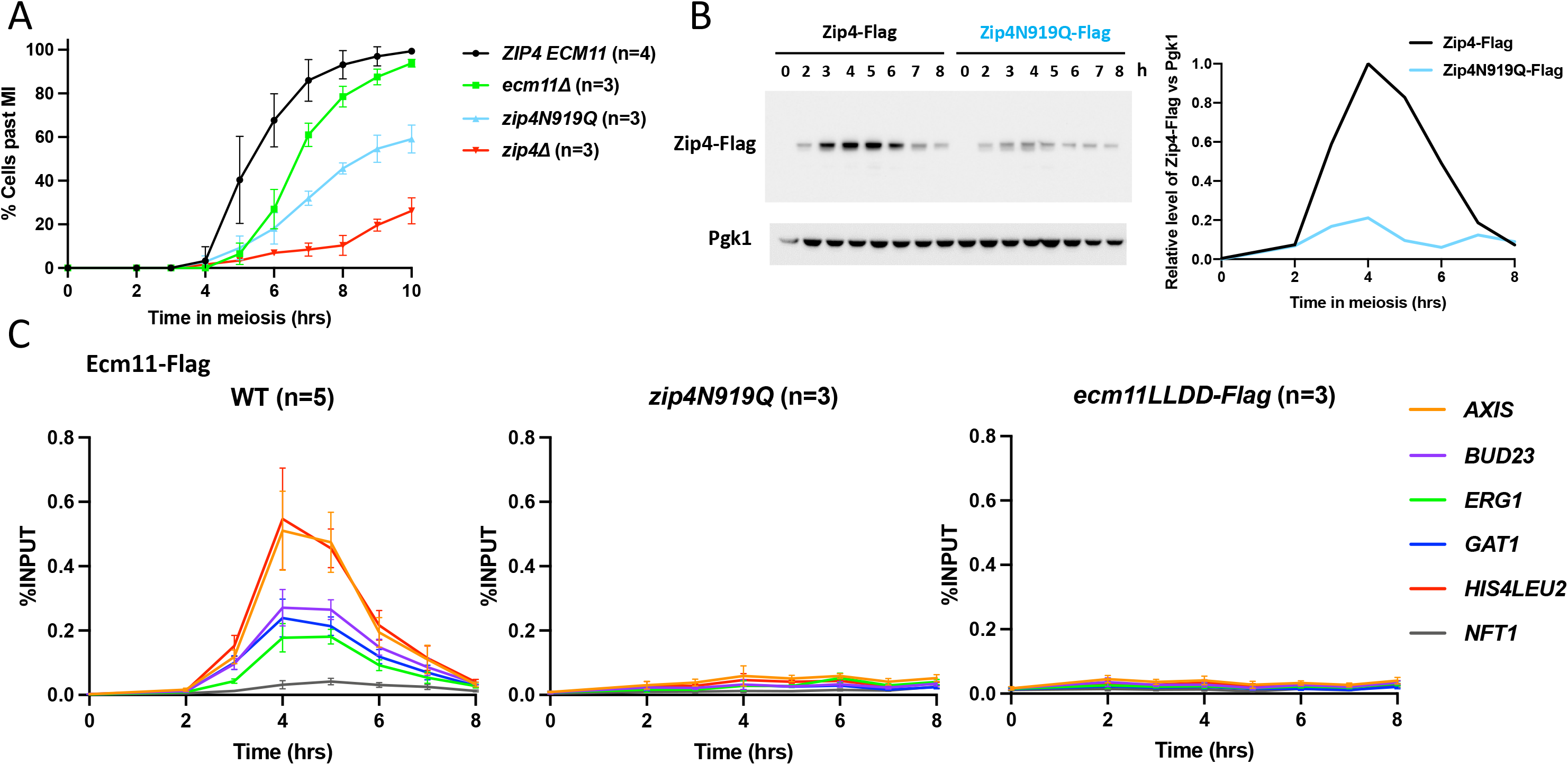
z*i*p4N919Q and *ecm11LLDD* phenotype in meiosis. A. Meiotic progression assessed by DAPI-staining of the strains with the indicated genotype. Values are the mean ± SEM of at least three independent experiments (except for ecm11LLDD-Myc: ±SD from two independent experiments). B. Western blot time course analysis of Zip4-Flag in wild-type cells or *ecm11*Δ strain, and Zip4N919Q-Flag. Right: quantification of Zip4-Flag signal, relative to Pgk1. C. ChIP monitoring of Ecm11-Flag in the indicated strains and Ecm11LLDD-Flag association with different chromosomal regions, measured by qPCR using primers that cover the indicated regions. Values are the mean ± SEM of at least three independent experiments.

**Figure S5:**
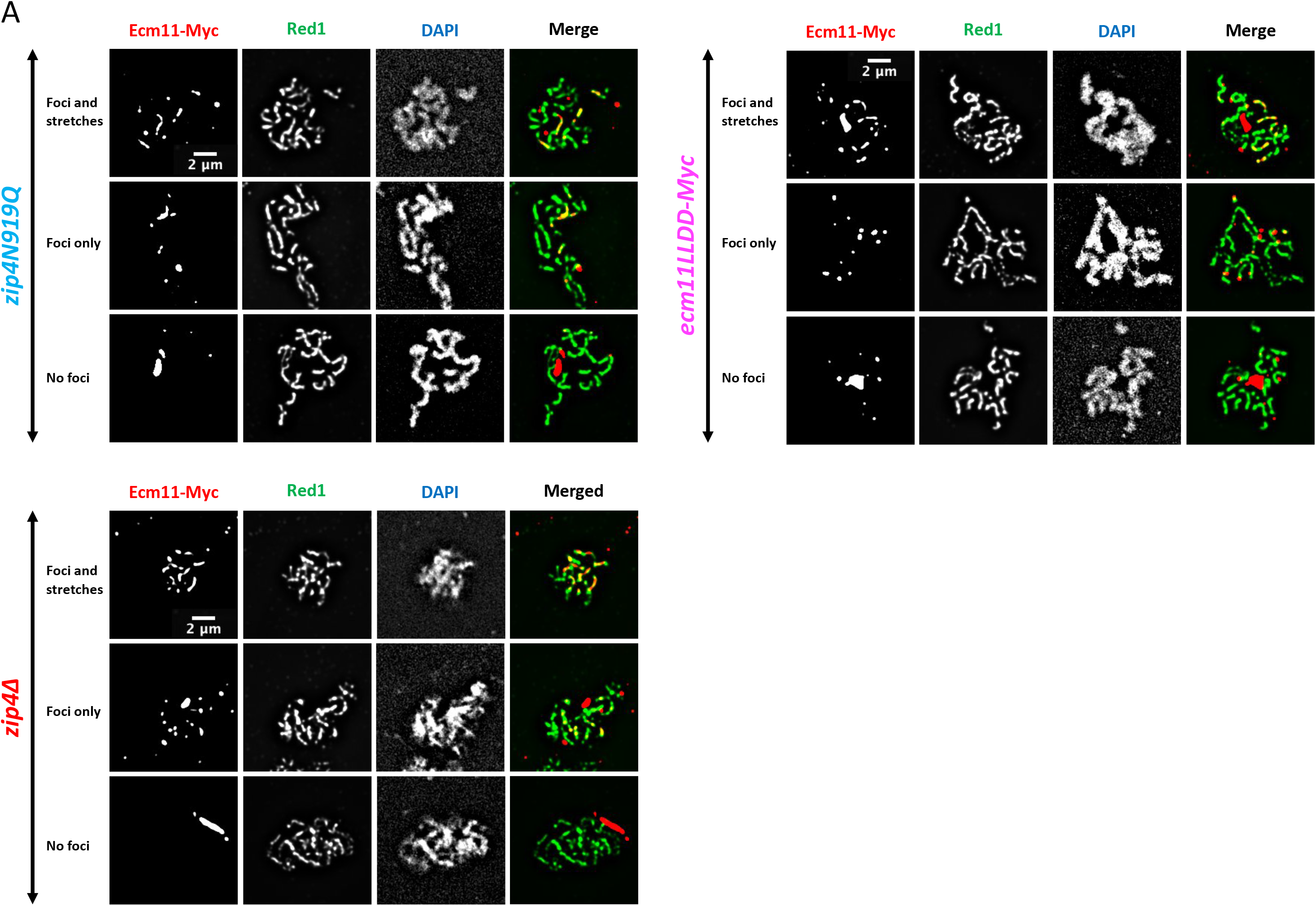
Localization of Ecm11-Myc on meiotic spreads. A. Ecm11-Myc localization on surface-spread chromosomes in the indicated strains. Red: anti-Myc; green: anti-Red1; blue: DAPI. The description of the categories is in Fig. 3C

**Figure S6:**
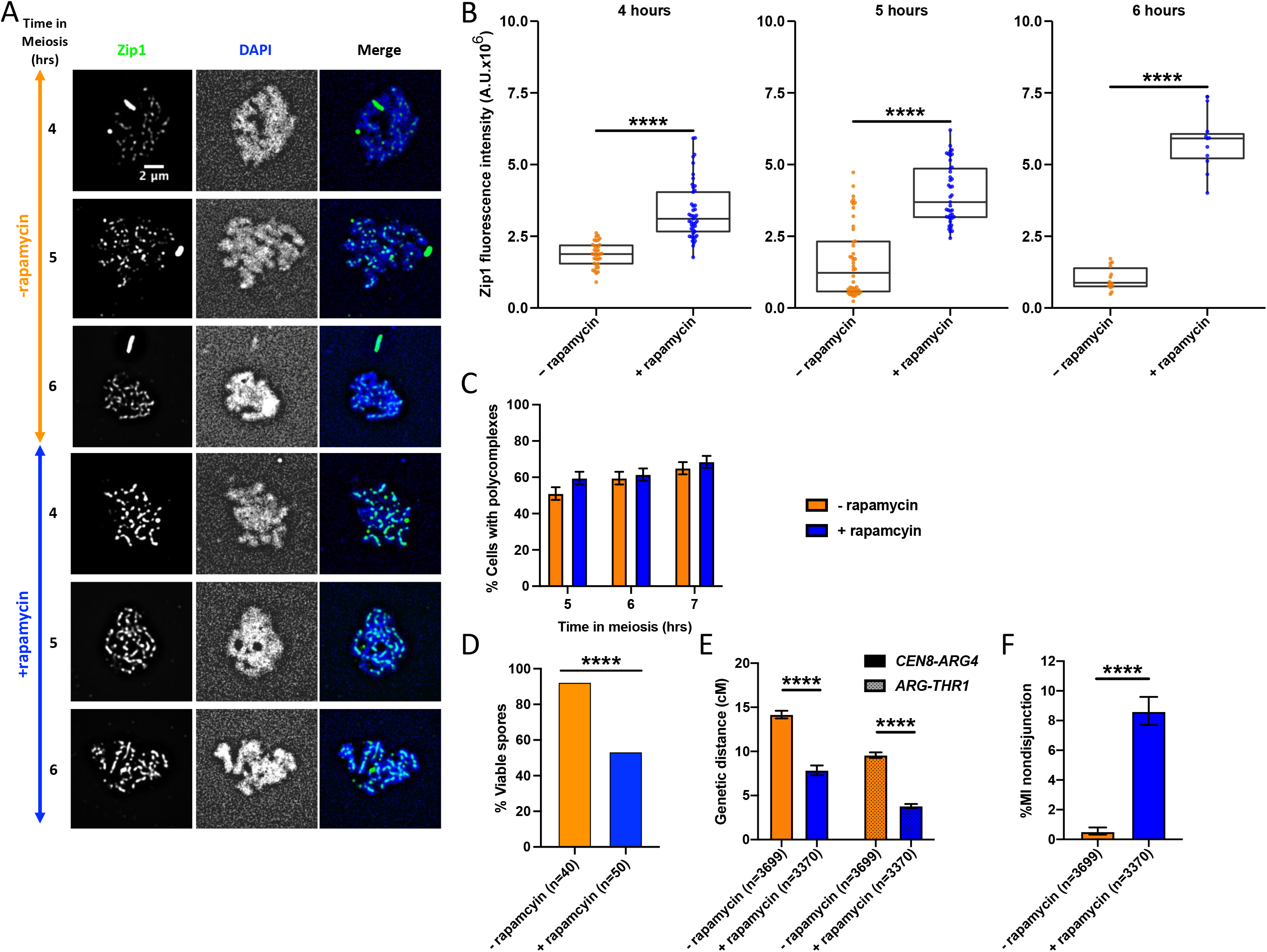
Zip1 staining and meiotic recombination after tethering Ecm11LLDD to Zip4. A. Zip1 localization on surface-spread chromosomes in the indicated conditions (- and + rapamycin) at 4, 5 and 6 hours after meiosis induction. Only pachytene or pachytene-like stages are considered. Green: anti-Zip1; blue: DAPI. B. Quantification of Zip1 signal intensity observed in A. ****: p-value<0.0001, Wilcoxon test. C. Quantification of DAPI-positive spreads showing a polycomplex. At least 200 spreads were considered for each condition. Values are % cells ± SD of the proportion. D. Spore viability in the indicated conditions (- and + rapamycin) 72 h after meiosis induction E-F. Crossing-over frequency and MI non disjunction. Same experimental setup as in Fig. 4. Genetic distances in the two genetic intervals *CEN8-ARG4* and *ARG4-THR1* on chromosome VIII are plotted as cM ± SE for the indicated genotypes. **\*\*\***\*: p- value<0.0001, G-test.

**Figure S7:**
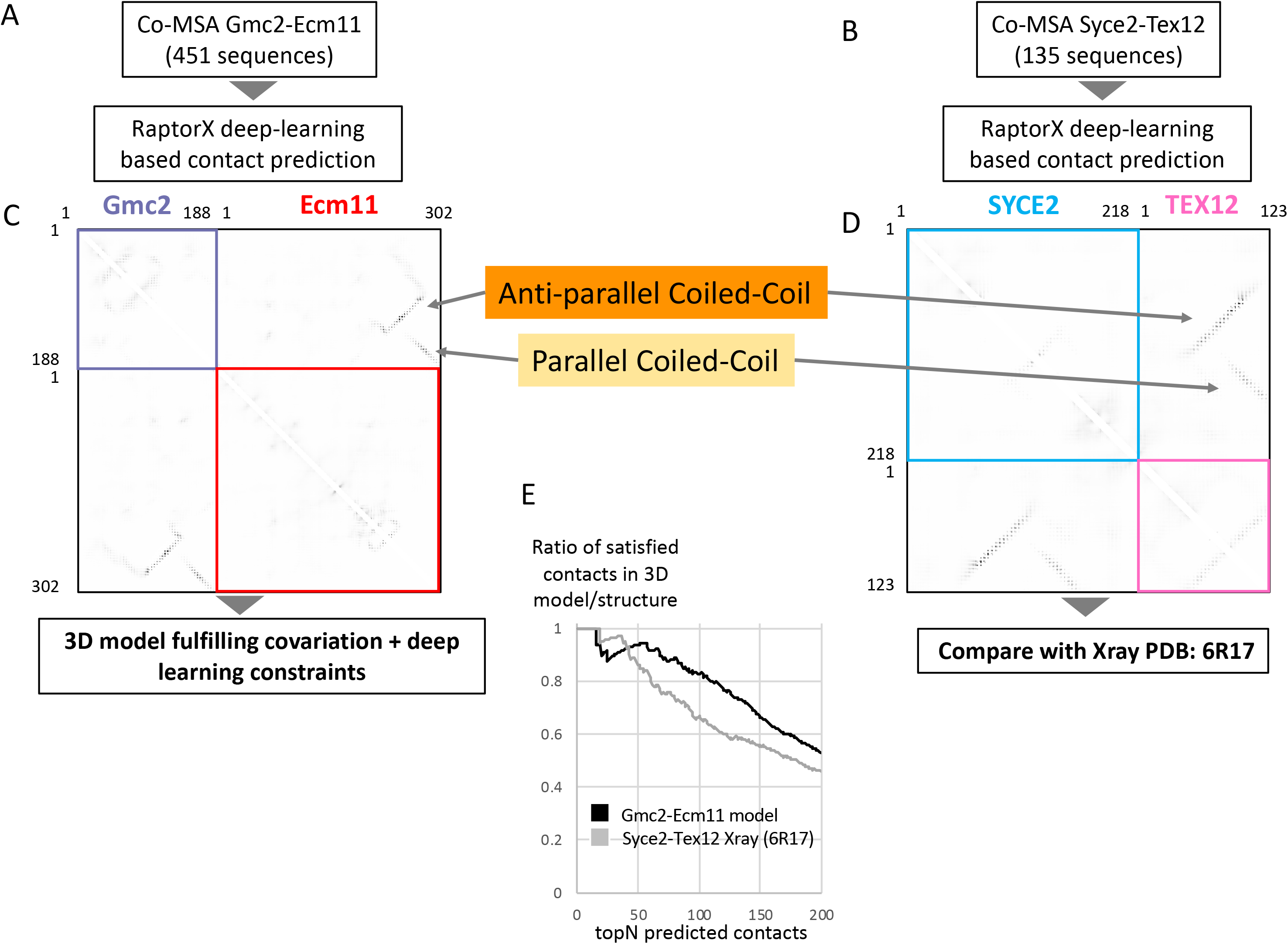
Modelling the assembly of the Gmc2-Ecm11 complex using constraints of deep learning-enhanced covariation-based prediction methods reveals similarities with the assembly of the TEX12-SYCE2 hetero-tetrameric coiled-coils. A. A co-multiple sequence alignment (co-MSA) containing 451 non-redundant pairs of fungal sequences homologous to *S. cerevisiae* Gmc2 and Ecm11 was concatenated and used as input of RaptorX contact prediction method (see STAR Methods). B. The same protocol was performed with homologs of human SYCE2 and TEX12 using a 135-sequences co-MSA. C. The contact maps predicted by RaptorX for Gmc2-Ecm11 are shown with a grey-scale representing contacts probabilities. Coloured boxes indicate the predicted intra-molecular contacts while the contacts outside the coloured boxes report the inter-molecular predicted contacts. Inter-molecular contacts are predicted significantly stronger with co-existence of anti-parallel (orange) and parallel (light orange) coiled-coils. RaptorX constraints with succession of anti-parallel and parallel coiled-coils could only be respected assuming a dimer of heterodimer for Gmc2-Ecm11 subunits, to build the 3D model (See Fig. 6D). D. The same predictions were also run to predict the contact map for the SYCE2-TEX12 complex. E. Analysis of the consistency between the contacts maps predicted using RaptorX and the 3D model of the Gmc2-Ecm11 complex shown in Fig. 6D (black curve) or the structure of the SYCE2-TEX12 complex shown in Fig. 6C (grey curve). The curves report the ratio of satisfied contacts between residues (distance Cb-Cb < 8Å) among the top N predicted contacts sorted by decreasing probabilities. The plots span the best 200 contacts. For both the crystal structure of SYCE2-TEX12 complex and the model of Gmc2-Ecm11, we observe that about 90% of the top50 predicted contacts are correct or can be satisfied, respectively. This comparison establishes the likelihood of the proposed assembly mode for the Gmc2-Ecm11 complex.

## Notes

### Competing Interest Statement

The authors have declared no competing interest.

